# Coordinated translational control of multiple immune checkpoints by the integrated stress response pathway in lung cancer

**DOI:** 10.1101/2024.10.23.619897

**Authors:** Shayna Thomas-Jardin, Shruthy Suresh, Ariana Arce, Nicole Novaresi, Emily Stein, Lisa Thomas, Cheryl Lewis, Chul Ahn, Bret M. Evers, Maria E. Salvatierra, Wei Lui, Khaja Khan, Luisa Maris Solis Soto, Ignacio Wistuba, John D. Minna, Kathryn A. O’Donnell

## Abstract

The integrated stress response (ISR) is an adaptive pathway hijacked by cancer cells to survive cellular stresses in the tumor microenvironment. ISR activation potently induces Programmed Death Ligand 1 (PD-L1), leading to suppression of anti-tumor immunity. Here we sought to uncover additional immune checkpoint proteins regulated by the ISR to elucidate mechanisms of tumor immune escape. We show that CD155 and PD-L1 are coordinately induced by the ISR, enhancing translation of both immune checkpoint proteins through bypass of inhibitory upstream open reading frames (uORFs) in their 5′ UTRs. Analysis of primary human lung tumors identifies a significant correlation between PD-L1 and CD155 expression. ISR activation accelerates tumorigenesis and inhibits T cell function, effects that can be overcome by combining PD-1 blockade with the ISR inhibitor ISRIB. These studies uncover a novel mechanism by which two immune checkpoint proteins are coordinately regulated and suggest a new therapeutic strategy for lung cancer patients.

**Statement of Significance:** This study uncovers a novel mechanism for the coordinated translational regulation of the PD- L1/PD1 and CD155/TIGIT immune checkpoint pathways and highlights the ISR as a therapeutic vulnerability for lung cancer. Inhibition of the ISR pathway bolsters PD-1 blockade, potentially unveiling a new therapeutic strategy for lung cancer patients.

## Introduction

The discovery of immune checkpoint pathways, and the development of clinically available checkpoint inhibitors, has dramatically improved therapeutic outcomes in a wide range of human malignancies. One critical immune checkpoint axis consists of programmed cell death protein 1 (PD-1), an inhibitory receptor expressed on T cells, and its binding partner programmed cell death ligand 1 (PD-L1) (1), expressed on tumor cells and other antigen presenting cells. Engagement of PD-1 with PD-L1 leads to suppression of T cell growth, survival, and other effector functions (2, 3). Clinically approved antibodies targeting these proteins restore T cell-mediated antitumor immunity, resulting in remarkable benefits for non-small cell lung cancer (NSCLC), melanoma, and kidney cancer patients (4, 5). Since its approval in 2015, PD-L1/PD-1 immune checkpoint blockade (ICB) has become a first-line therapy for lung cancer patients (5–7). Despite this remarkable progress, less than 30% of NSCLC patients respond to ICB with durable benefit. Thus, there is a growing need to identify mechanisms of resistance to ICB and to discover additional immune checkpoint therapies that may be combined with PD-L1/PD-1 blockade to improve therapeutic responses.

To elucidate the mechanisms through which PD-L1 is upregulated in human lung cancer cells, we previously performed a genome-wide loss-of-function CRISPR/Cas9 screen to identify novel regulators of PD-L1 (8). This led to the unexpected finding that *PD-L1* can be induced at the level of translation. The most significant hit among the negative regulators of PD-L1 was *Uroporphyrinogen Decarboxylase (UROD*), a key enzyme in the heme biosynthesis pathway. Impairment of heme production by genetic depletion of *UROD* or chemical inhibition of the heme pathway potently induced PD-L1 protein, without a corresponding increase in *PD-L1* mRNA levels (8). Disruption of heme synthesis is known to activate the cellular stress response pathway known as the integrated stress response (ISR) (9, 10). We demonstrated that ISR activation enhances translation of *PD-L1*, through a mechanism involving the bypass of inhibitory upstream open reading frames (uORFs) in the *PD-L1* 5′-untranslated regions (UTRs). Furthermore, we found that *Urod* depletion accelerates tumorigenesis by suppressing CD8+ T- cells. Moreover, the immunosuppressive effects of *Urod* inhibition are dependent upon the PD- 1/PD-L1 axis, thereby supporting a role for the ISR in enhancing tumor growth through suppression of anti-tumor immune responses.

Eukaryotic cells activate the ISR to cope with diverse cellular stresses, to restore homeostasis, or initiate cell death (10–12). These stresses activate one of four kinases, respectively: ER stress and hypoxia activate PKR-like ER kinase (PERK), viral infection activates protein kinase double-stranded RNA-dependent (PKR), amino acid deprivation activates general control non-derepressible-2 (GCN2), and heme deprivation activates heme regulated inhibitor (HRI) (9, 10, 12). Upon activation, each of these kinases phosphorylates the alpha subunit of eukaryotic translation initiation factor 2 alpha (eIF2α) at serine 51 (9, 10). This inhibits the guanine nucleotide exchange activity of eIF2B by forming a sequestered eIF2-eIF2B complex, leading to impaired eIF2 recycling. As a result, phosphorylation of eIF2α limits the formation of the ternary complex required for translation initiation (eIF2α:GTP:Met-tRNA), thereby protecting cells by attenuating global protein translation. Paradoxically, specific messenger RNAs (mRNAs) encoding proteins that are critical for relieving cellular stress are translated more efficiently upon activation of the ISR (9, 10, 13). These stresses, especially oxidative stress, nutrient deprivation, and hypoxia, are common in the tumor microenvironment (TME). Thus, cancer cells exploit the ISR to enhance survival in the harsh setting of the TME.

There is a growing appreciation that ISR activation promotes tumor progression and immune evasion (12). Increased eIF2α phosphorylation is commonly observed in human lung tumors (14), and correlates with poor clinical outcome of lung adenocarcinoma patients (15). ISR activation enhances tumorigenesis through multiple mechanisms, including translation of oncogenic transcripts (16), increased invasion and metastasis (17), and increased cell survival (18). Both human and mouse PD-L1 harbor inhibitory uORFs in their 5′ UTRs that suppress baseline translation of PD-L1 at the canonical AUG. Transgenic MYC expression activates the ISR to overcome uORF-mediated inhibition and drive *Pd-l1* translation in a liver cancer mouse model (19). Similarly, ISR activation through heme deficiency allows the bypassing of inhibitory uORFs and enhances *PD-L1* translation in human lung cancer cells. Thus, translational control of the PD-L1 immune checkpoint under physiologic or oncogenic stress represents an important mechanism of immune evasion in human cancers.

In the current study, we sought to identify additional immune checkpoint proteins that are regulated by the ISR pathway and elucidate shared mechanisms of tumor immune escape in human lung cancers. We activated the ISR pathway and screened for induction of key immune checkpoint proteins known to be expressed in human lung cancers. We discovered that Cluster of differentiation 155 (CD155), also known as the Poliovirus receptor (PVR), and PD-L1 are coordinately induced by the ISR pathway, leading to enhanced translation of both immune checkpoint proteins through bypass of inhibitory uORFs in their 5′ UTRs. We found a significant correlation between CD155 and PD-L1 in a large panel of primary human lung tumors, and we further demonstrated that ISR activation promotes tumorigenesis and inhibits T cell function in co-culture assays and in mouse syngeneic models. Moreover, inhibition of the ISR pathway with the small molecule, Integrated Stress Response Inhibitor (ISRIB), enhanced the response to PD-1 blockade *in vivo*. Collectively, our studies illuminate a novel mechanism by which two critical immune checkpoint proteins are coordinately regulated and suggest a new therapeutic strategy for lung cancer patients.

## Results

### ISR pathway activation induces CD155 and PD-L1 protein

Based on our finding that ISR activation drives *PD-L1* translation (8), we hypothesized that additional immune checkpoint proteins may be coordinately regulated by the ISR in NSCLC, potentially facilitating resistance to anti-PD-1/PD-L1 therapies. To test this hypothesis, we activated the ISR by thapsigargin treatment, an inducer of ER stress, in multiple independent *KRAS* mutant (H441, H358) and *EGFR* mutant (PC9, HCC827) NSCLC cell lines and screened for elevated expression of key immune checkpoint proteins that are expressed in human lung cancers (20–26). These included PD-L2, CD155, Galectin-3, Galectin-9, and HVEM, which are all capable of inhibiting immune cell function through binding to associated receptors on T cells and other immune cell subtypes (**Supplementary Fig. S1A**). Among these factors, we observed that CD155 was induced by thapsigargin treatment in all tested cell lines (**Supplementary Fig. S1B**).

CD155 is a cell surface protein that engages with several immune cell receptors to facilitate immune suppression, including T Cell Immunoreceptor with Ig and ITIM Domains (TIGIT), which is expressed on T cells, dendritic cells, and NK cells (25–29). In addition to this role in immune suppression, CD155 participates in cell adhesion and motility (30, 31), and is overexpressed in multiple human malignancies including lung adenocarcinoma (23, 26). We found that CD155 and PD-L1 are coordinately induced by Salubrinal, a selective inhibitor of eIF2α de-phosphorylation (32) (**Fig. 1A**). Additionally, amino acid starvation (**Fig. 1B**) and ER stress stimulated by Thapsigargin treatment (**Fig. 1C**) induced both immune checkpoint proteins. We previously demonstrated that depletion of UROD, an essential enzyme in the heme biosynthesis pathway, induces the ISR and, consequently, PD-L1 levels through disruption of heme availability (8). Accordingly, CRISPR/Cas9 knockout of *UROD* (**Fig. 1E**), or transient knockdown with siRNA (**Fig. 1F**), induced both CD155 and PD-L1 without a corresponding increase in *CD155* or *PD-L1* mRNA (**Supplementary Fig. S1C**). Similarly, *CD155* or *PD-L1* mRNA levels were not induced by Salubrinal treatment or amino acid deprivation (**Supplementary Fig. S1D-E**), consistent with ISR-mediated translational regulation. As expected, *DNA damage-inducible protein 34 (GADD34)*, a target of the key ISR effector activating transcription factor 4 (ATF4), was induced under these conditions. Moreover, flow cytometry confirmed elevated cell surface expression of CD155 and PD-L1 protein (**Fig. 1D**), indicating functional immune checkpoint expression capable of engaging immune cells.

**Figure 1.**
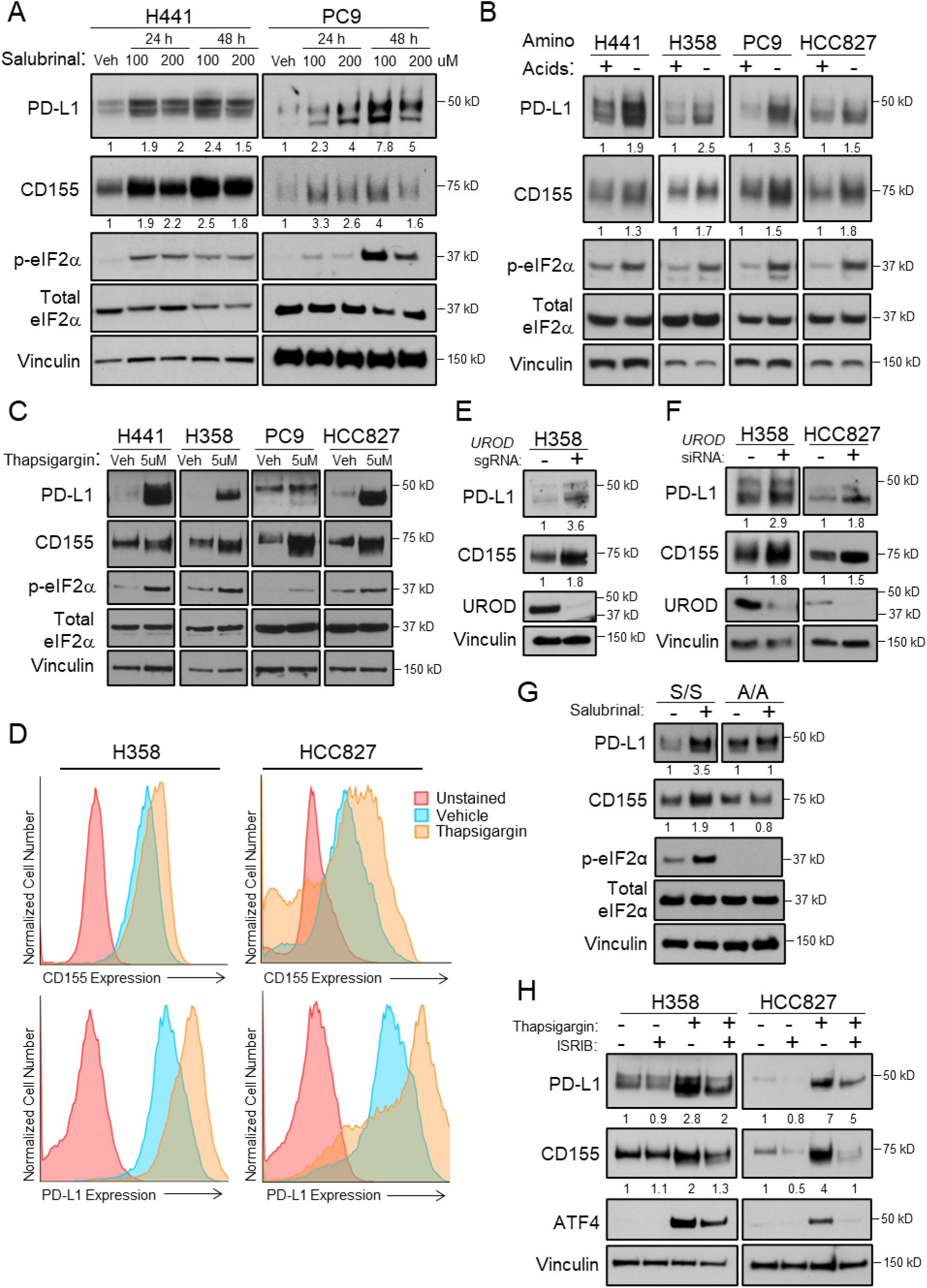
ISR pathway activation induces PD-L1 and CD155 protein. (**A**) Western blot analysis of human *KRAS* mutant H441 or *EGFR* mutant PC9 cells treated for 24 or 48 hours with 100uM or 200uM Salubrinal or DMSO vehicle control. Vinculin served as a loading control for this and subsequent western blots. (**B**) Western blot analysis of human H441, H358, PC9, and HCC827 cells after 24 hours in RPMI 1640 media supplemented with or without amino acids. (**C**) Western blot analysis of human H441, H358, PC9, and HCC827 cells treated for 24 hours with 5uM thapsigargin (ER Ca+ ATPase pump inhibitor) to induce ER stress, or DMSO vehicle control. (**D**) Flow cytometric analysis of cell-surface CD155 and PD-L1 protein in DMSO vehicle control and thapsigargin-treated (5uM for 24 hours) H358 or HCC827 NSCLC cells. (**E**) Western blot analysis of H358 cells with control or *UROD* sgRNA. (**F**) Western blot analysis of H358 or HCC827 cells with control or *UROD* siRNA. (**G**) Western blot analysis in eIF2⍺ wildtype (S/S) or mutant (Ser51Ala A/A) mouse embryonic fibroblasts (MEFs) treated with DMSO vehicle control or 100uM Salubrinal for 24 hours. (**H**) Western blot analysis of thapsigargin and ISRIB treated human NSCLC cells. Data from a single experiment are shown and are representative of at least 3 independent experiments.

To determine whether a functional ISR pathway is necessary for increased expression of both immune checkpoint proteins, we treated mouse embryonic fibroblasts (MEFs) expressing either wildtype eIF2α (S/S cells) or mutant eIF2α with serine-51 mutated to alanine (A/A cells) with Salubrinal. ISR activation led to phosphorylation of eIF2α at serine-51 and induction of CD155 and PD-L1 in S/S cells, but not in A/A cells (**Fig. 1G**). Conversely, treatment with the small molecule inhibitor of the ISR pathway, ISRIB (33, 34), diminished CD155 and PD-L1 levels in thapsigargin treated *KRAS* and *EGFR* mutant NSCLC cells (**Fig. 1H**). These data demonstrated that both PD-L1 and CD155 are induced by the ISR in human lung cancer cells.

### ISR activation enhances CD155 translation

To test if CD155 and PD-L1 are selectively translated during ISR activation, we performed polysome profiling by sucrose gradient ultracentrifugation of control and Salubrinal treated cells. ISR activation resulted in an overall reduction in polysome abundance, indicative of decreased global translation (**Fig. 2A**). As expected, *PD-L1* and *ATF4* mRNA redistributed to heavier polysome fractions upon ISR activation (**Fig. 2B, Supplementary Fig. S2A-C**). Analysis of *CD155* mRNA in pooled or individual polysome fractions demonstrated that it similarly redistributed to heavier polysomes after Salubrinal treatment, indicating increased translation (**Fig. 2C, Supplementary Fig. S2D**). This effect was confirmed using a second independent primer set (**Supplementary Fig. S2E, S2F)**. Analysis of Actinomycin D-treated and cycloheximide-treated H441 cells confirmed that the increased protein abundance and redistribution of mRNA to heavier polysomes is not due to enhanced mRNA or protein stability, respectively (**Supplementary Fig. S2G-H**).

**Figure 2.**
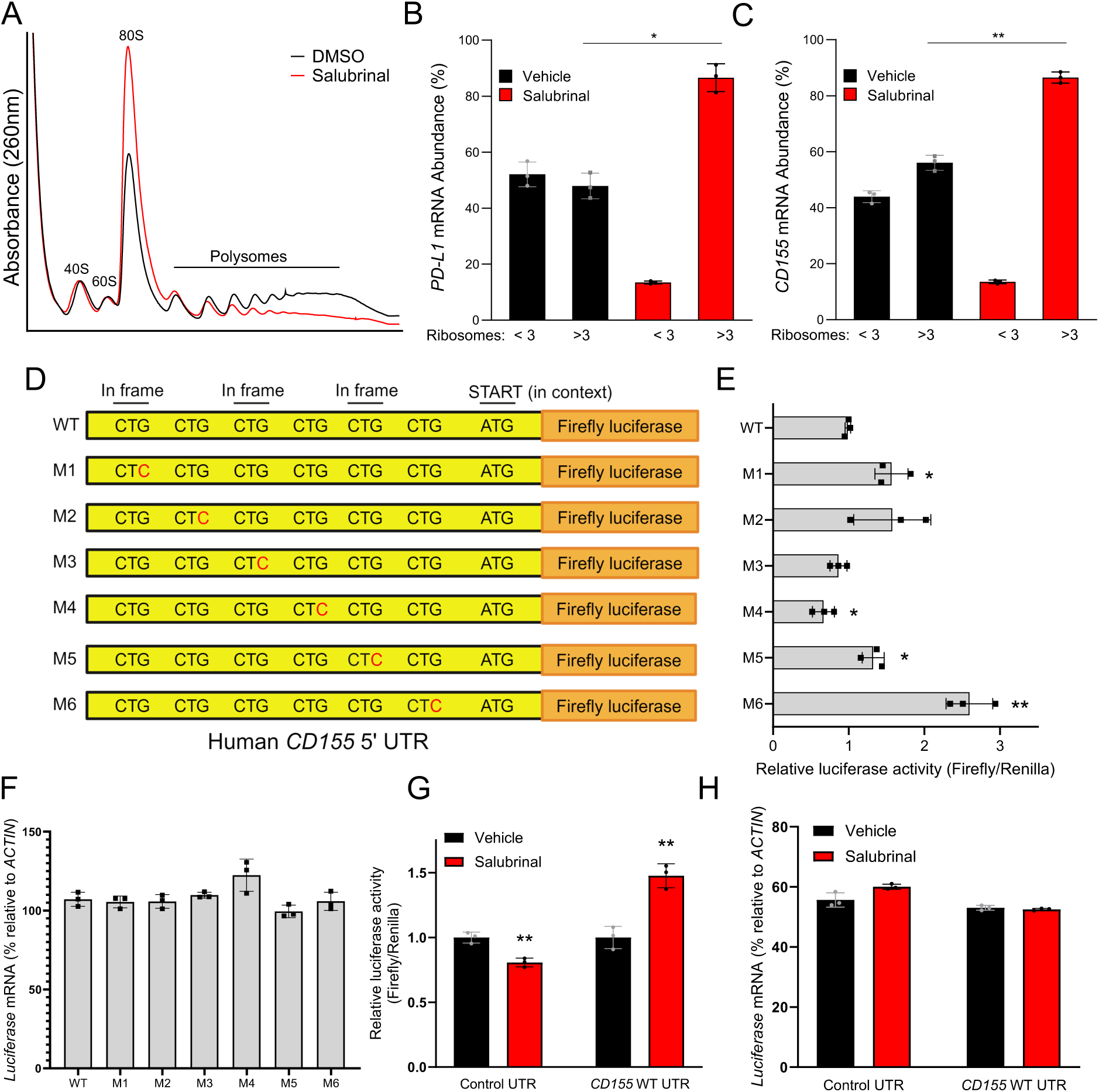
ISR pathway activation enhances *PD-L1* and *CD155* translation. (**A**) Polysome profiling of H1944 Vehicle or Salubrinal-treated cells (100uM for 24 hours). (**B**) Quantitative real-time PCR analysis of *PD-L1(CD274)* and *CD155(PVR)* mRNA (**C**) in ribosomal fractions from (**A**). qRT-PCR analysis for each gene shown was performed with 1 primer set spanning an exon-exon junction (Primer Set 1). Data for Primers Set 2 is available in Supplementary Figure 2. Fractions associated with <3 ribosomes were grouped to represent poorly translated mRNAs, fractions associated with >3 ribosomes were grouped as efficiently translated mRNAs. *PD-L1* and *CD155* mRNA expression in each fraction was normalized to *Luciferase* and mRNA abundance was calculated as the % of total in all fractions. Luciferase control mRNA was added to each fraction prior to RNA extraction to control for variability. Error bars represent standard deviation from the mean from three independent fractions (<3 or >3 ribosomes). (**D**) Diagram of the wildtype human *CD155* 5′ UTR with 6 CTGs, and mutant constructs with CTGs mutated to CTCs, cloned upstream of a luciferase reporter. (**E**) Dual luciferase assay of MEFs transfected with indicated *CD155*-5′ UTR-Firefly luciferase reporter constructs normalized to co-transfected control Renilla luciferase. Luciferase activity was monitored after 48h. Error bars represent standard deviation from the mean from n=3 biological replicates. Data from a single experiment are shown and representative of three independent experiments. (**F**) qRT-PCR analysis of mean luciferase mRNA normalized to actin in MEFs shown in (**E**). Error bars represent standard deviation from the mean from n=3 biological replicates. (**G**) Dual luciferase assay of the *CD155* 5′ UTR in MEFs, Vehicle or Salubrinal-treated (100uM for 24 hours). Error bars represent standard deviation from the mean from n=3 biological replicates. (**H**) qRT-PCR analysis of mean luciferase mRNA normalized to actin in MEFs shown in (**G**). n=3 biological replicates. A Student’s t-test was used to determine statistical significance (* p<0.05, ** p<0.005, *** p<0.0005).

Mechanistically, ISR activation has been shown to induce the translation of specific mRNAs that harbor inhibitory upstream open reading frames (uORFs), including canonical ISR response transcripts such as *ATF4*, *GADD34*, and *GCN4* (16, 35, 36). The phosphorylation of eIF2α is hypothesized to weaken its ability to form an active ternary complex (eIF2α:GTP:Met-tRNA), thereby leading to leaky scanning that can bypass inhibitory uORFs in the 5′ UTR and increase translation at canonical translation start sites. We and others (8, 19) previously found that the *PD-L1* 5′ UTR contains 5 uORFs initiating with the non-canonical start codon CUG, and that mutation of the third, fourth, or fifth uORF leads to an increase in translation of a reporter construct (8), suggesting that activation of the ISR promotes bypass of these inhibitory uORFs, enhancing PD-L1 translation. To investigate the contribution of uORFs to CD155 translation, we examined the human *CD155* 5′ UTR and identified 6 upstream CUGs (3 out-of-frame, 3 in-frame within 175 bp upstream of the annotated AUG start codon). We generated constructs to test the effect of these uORFs on CD155 translation by cloning the wild-type 5′ UTR, or mutant versions with each CUG mutated to CUC, upstream of firefly luciferase (**Fig. 2D**). These constructs were transfected into MEFs along with a control Renilla luciferase reporter. Mutation of the first, fifth, and most significantly the sixth uORF enhanced firefly luciferase activity (**Fig. 2E**), without affecting luciferase mRNA levels (**Fig. 2F**). Similarly, ISR activation enhanced firefly luciferase activity of a wild-type CD155 5′ UTR reporter, but not a control UTR reporter without impacting luciferase mRNA abundance (**Fig. 2G**, **2H**). Collectively, these results support a model wherein ISR activation promotes the bypass of inhibitory uORFs in the CD155 and PD-L1 5′ UTRs, resulting in enhanced translation and immune checkpoint activation.

### ISR pathway activation diminishes immune cell function and infiltration and promotes tumorigenesis

We next sought to determine the extent to which activation of the ISR pathway in tumor cells impacts immune cell responses both *in vitro* and *in vivo*. Our prior data demonstrated that *Urod* knockdown accelerates tumor growth by suppressing CD8^+^ T-cells (8). Moreover, the immunosuppressive effects of *Urod* knockdown are dependent upon the PD-1/PD-L1 axis. Based on these data, we hypothesized that tumors undergoing ISR activation will express higher levels of PD-L1 and CD155 protein, which will engage their respective receptors on immune cells in the microenvironment, and thereby suppress immune cell proliferation and function. To evaluate the effects of ISR activation on the tumor-immune cell milieu, we first performed co-culture assays with H358 NSCLC cells and Jurkat T cells, and then assessed T cell activation and cytotoxic activity with IL-2 and Granzyme-B ELISA assays from conditioned media. Treatment of H358 cells with Salubrinal resulted in reduced secretion of IL-2, a marker of T cell activation, and Granzyme B, a marker of cytotoxic T cell function, from co-cultured Jurkat cells (**Fig. 3A**). IL-2 and Granzyme B levels were significantly rescued by ISRIB treatment in co-culture assays. Similarly, *UROD* knockout in H358 cells reduced levels of IL-2 and Granzyme B secretion from co-cultured Jurkat cells, an effect also significantly rescued by ISRIB treatment (**Fig. 3B**). We observed similar results in co-cultures of H358 and Jurkat cells following amino acid starvation (**Supplementary Fig. S3A**).

**Figure 3.**
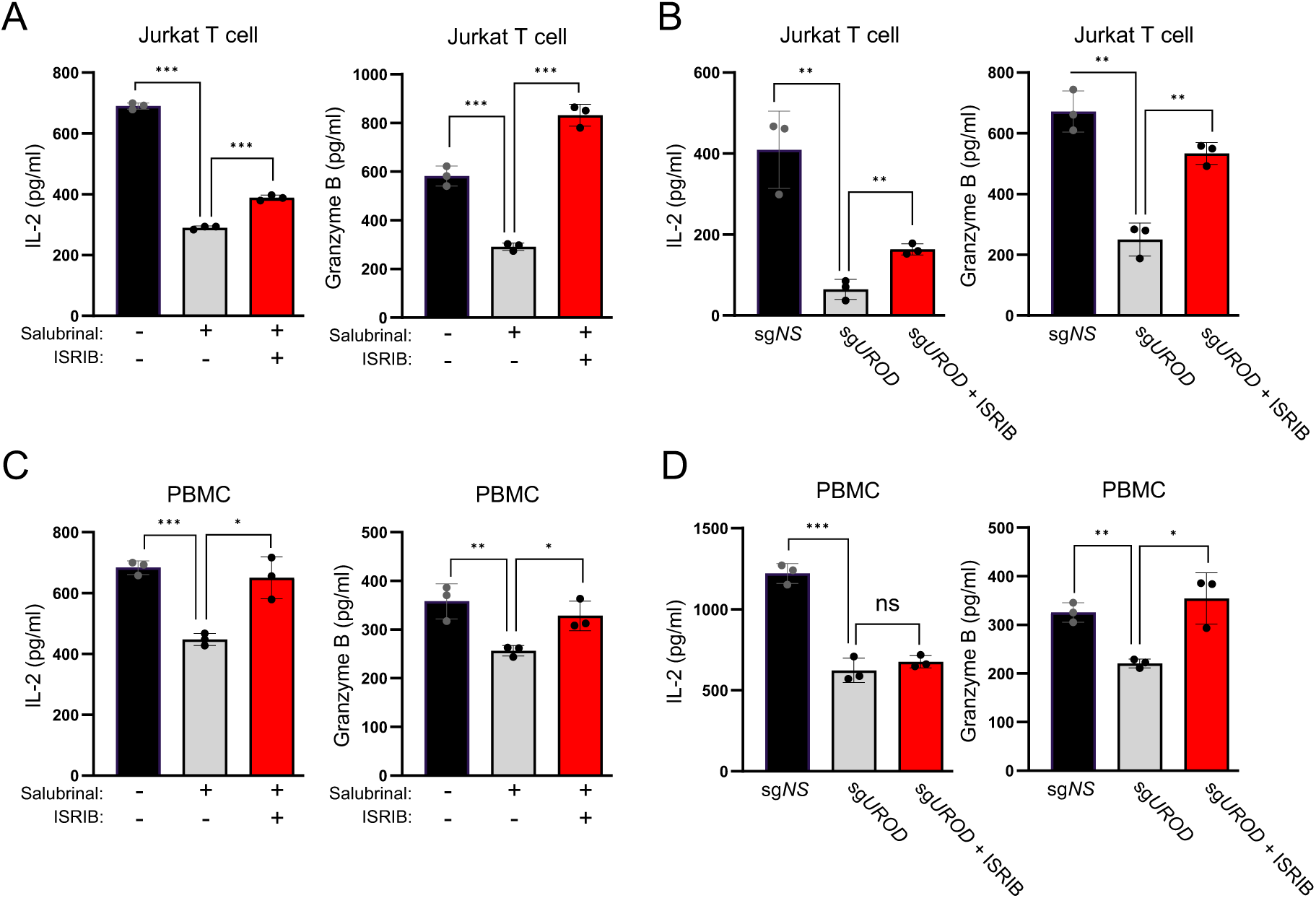
ISR pathway activation diminishes immune cell function *in vitro*. (**A**) ELISAs for IL-2 and Granzyme B of Jurkat T cells co-cultured with H358 cells. H358 cells were pre-treated for 24 hours with Salubrinal and 800nM ISRIB, then washed and co-cultured with Jurkat T cells for an additional 24 hours with α-CD3 and α-CD28 activating antibodies (4ug/ml each). Error bars represent standard deviation from the mean from n=3 biological replicates. Data from a single experiment are shown and representative of three independent experiments. (**B**) ELISAs for IL-2 and Granzyme B of Jurkat T cells co-cultured with H358 cells with control or *UROD* sgRNA and 800nM ISRIB. H358 cells were co-cultured with Jurkat T cells for 24 hours with α-CD3 and α-CD28 activating antibodies (4ug/ml each). Error bars represent standard deviation from the mean from n=3 biological replicates. Data from a single experiment are shown and representative of two independent experiments. (**C**) ELISAs for IL-2 and Granzyme B of primary human PBMCs co-cultured with H358 cells. H358 cells were pre-treated for 24 hours with Salubrinal and 800nM ISRIB, then washed and co-cultured with PBMCs for an additional 24 hours with α-CD3 and α-CD28 activating antibodies (1ug/ml each). Error bars represent standard deviation from the mean from n=3 biological replicates. Data from a single experiment are shown and representative of three independent experiments. (**D**) ELISAs for IL-2 and Granzyme B of PBMCs co-cultured with H358 cells with control or *UROD* sgRNA and 800nM ISRIB. H358 cells were co-cultured with PBMCs for 24 hours with α-CD3 and α-CD28 activating antibodies (1ug/ml each). Error bars represent standard deviation from the mean from n=3 biological replicates. Data from a single experiment are shown and representative of two independent experiments. A Student’s t-test was used to determine statistical significance (* p<0.05, ** p<0.005, *** p<0.0005).

To investigate the effects of ISR activation in a more immunologically diverse system, we performed a similar series of experiments with H358 cells co-cultured with human peripheral blood mononuclear cells (PBMCs). Consistent with our results from H358-Jurkat co-culture experiments, we documented reduced IL-2 and Granzyme B secretion from PBMCs co-cultured with Salubrinal-treated or UROD-deficient H358 cells (**Fig. 3C-D**). Again, these effects were largely reversed by inhibition of the ISR with ISRIB.

We next investigated the effects of ISR activation directly on immune cell populations. The JAWS II murine dendritic cell line, primary mouse dendritic cells, and Jurkat T cells all have an intact ISR pathway, as evidenced by an increase in ATF4 protein upon activation of the ISR by multiple stressors (**Supplementary Fig. 3B-D**). Interestingly, however, we observed no detectable increase in PD-L1 or CD155 protein upon ISR activation in these cells. Thus, these immune checkpoint proteins appear to be refractory to ISR regulation in specific immune cell populations.

To evaluate the effects of ISR activation on tumorigenesis and immune infiltration *in vivo*, we utilized the CMT167 syngeneic lung cancer model, which harbors an oncogenic *Kras*^G12V^ mutation (37, 38). Furthermore, tumor growth can be suppressed in this model by PD-1 or PD- L1 antibody blockade (38). We confirmed that CD155 and PD-L1 are coordinately induced by Salubrinal in CMT167 cells (**Fig. 4A**) and by thapsigargin-induced ER stress (**Supplementary Fig. S4A**). CMT167 cells were then transplanted into immunocompetent C57BL/6 mice and daily Salubrinal or vehicle treatment was initiated. Salubrinal significantly enhanced tumor growth rates (**Fig. 4B**). Similarly, Salubrinal-treated tumors exhibited higher volume and weight at endpoint (**Fig. 4C**, **4D**). Fluorescent multiplex immunohistochemistry (IHC) analysis of tumors further revealed a significant reduction in total T cell infiltration, including CD3^+^, CD4^+^ and CD8^+^ T cells (**Fig. 4E-H**). Overall, these findings demonstrated that ISR activation enhances translation of *CD155* and *PD-L1*, promotes tumor growth, and suppresses host T cell infiltration *in vivo*.

**Figure 4:**
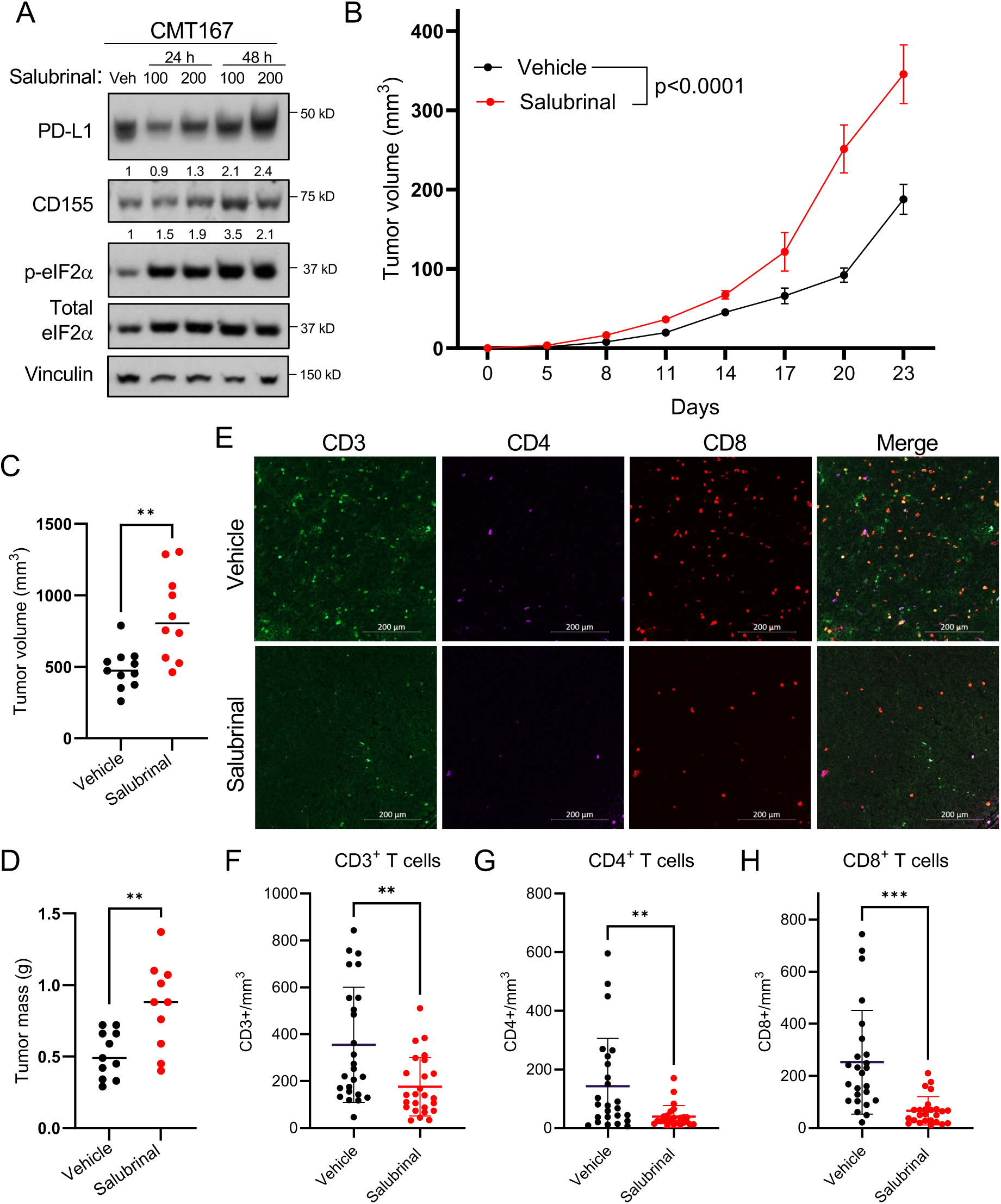
ISR activation enhances tumorigenesis and reduces immune cell infiltration *in vivo*. (**A**) Western blot analysis of *KRAS* mutant murine CMT167 cells treated for 24 or 48 hours with 100uM or 200uM Salubrinal or DMSO vehicle control. (**B**) Quantification of tumor volumes of CMT167 cells transplanted in C57BL/6J mice (n=11 mice per group for vehicle treated mice, n=10 mice per group for Salubrinal treated mice). Graph represents mean tumor volumes and error bars represent standard error of the mean. (**C**) End-point tumor volumes and (**D**) tumor mass of resected tumors shown in (**B**). Horizontal bars represent mean values. (**E**) Multiplexed IHC-F was performed on tumors from (**B**), 10X representative images shown, scale bar=200μm. (**F,G,H**) Quantification of CD3^+^, CD4^+^, and CD8^+^ tumor infiltrating lymphocytes (TILs, expressed as counts/mm^3^). 5 fields were quantified per mouse from 5 mice, graphs represent mean counts and error bars represent standard deviation from the mean. A Student’s t-test was used to determine statistical significance (* p<0.05, ** p<0.005, *** p<0.0005).

### Inhibition of the ISR pathway *in vivo* enhances response to PD-1 blockade

ISR inhibiting drugs ISRIB or PERK inhibitor GSK2606414 impair lung tumor formation in *KRAS* mutant xenografts and in syngeneic mouse models (15), however the impact of these drugs on anti-tumor immune responses has not been investigated. Given our data demonstrating that ISRIB treatment diminishes CD155 and PD-L1 induction in ISR-activated human NSCLC cells and in a syngeneic mouse model (**Fig. 1H, Supplementary Fig. S4A**), we next sought to evaluate the efficacy of combining ISR inhibition with immune checkpoint blockade *in vivo*. We hypothesized that restoration of eIF2-mediated protein translation with ISRIB treatment would reduce PD-L1 and CD155 protein levels, elicit an immune response, and synergize with PD-1 blockade. To test this hypothesis, we utilized doxycycline-inducible *Urod* shRNAs, which robustly induce CD155 and PD-L1 protein levels in CMT167 cells (**Fig. 5A**). CMT167 cells expressing *Urod* shRNA or *scrambled* shRNA control (**Fig. 5B-C**, **Supplementary Fig. S4B-C**) were subcutaneously transplanted into C57BL/6 mice and maintained on doxycycline water for the duration of the experiment. Mice in both groups were dosed daily with ISRIB or vehicle, while anti-PD-1 antibody or control IgG was administered every 3 days. As expected, *Urod* knockdown enhanced tumor growth *in vivo* (**Supplementary Fig. S4B**). ISRIB treatment modestly reduced tumor burden of *Urod* shRNA-treated tumors (**Fig. 5C, Supplementary Fig. 4C**). PD-1 blockade also reduced tumor burden of both *Urod* shRNA and *scrambled* shRNA control tumors, while the greatest effect was observed with the combination treatment of ISRIB and PD-1 blockade (**Fig. 5C).**

**Figure 5.**
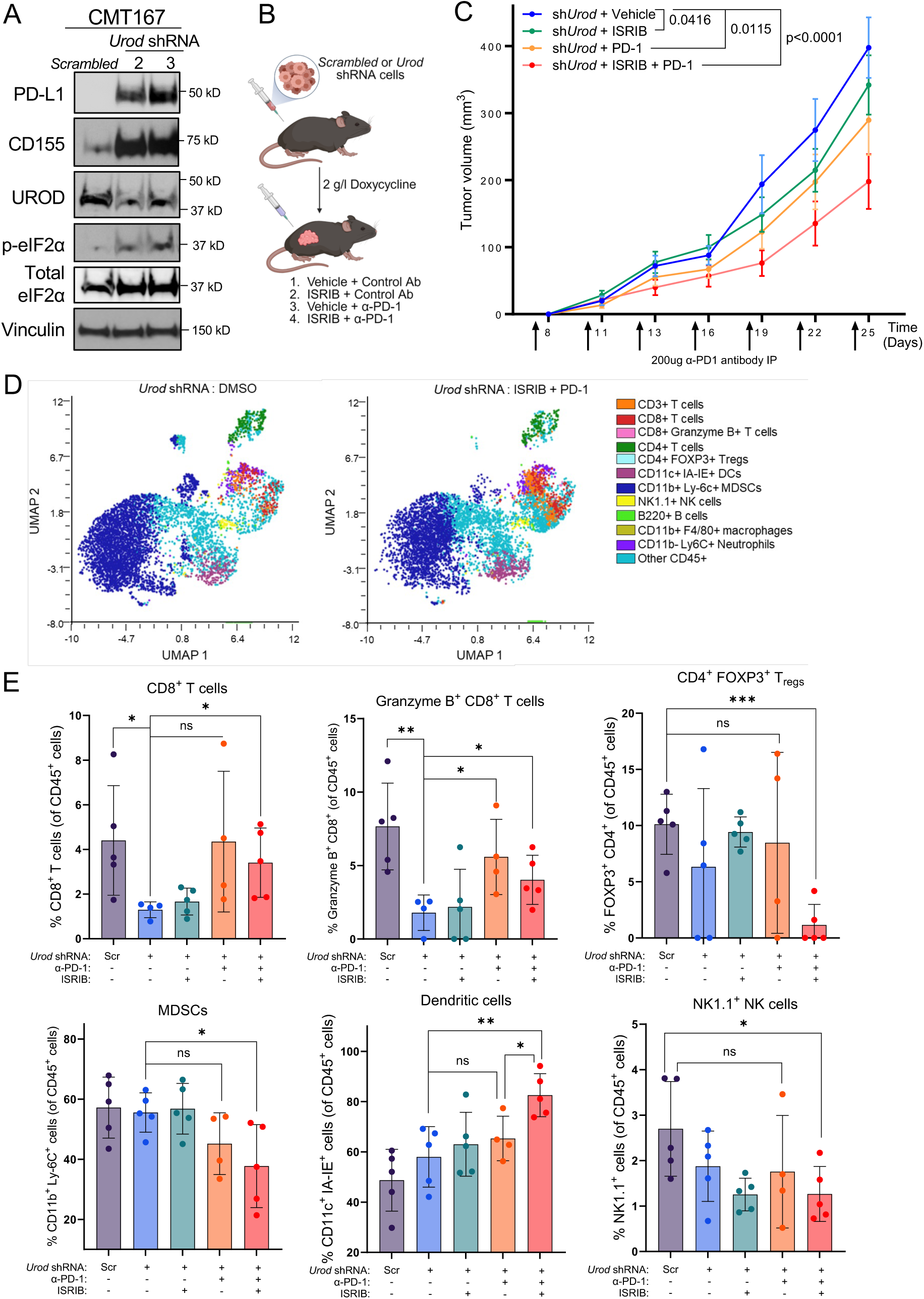
ISR pathway inhibition improves response to PD-1 blockade in a syngeneic mouse model. (**A**) Western blot analysis of *Kras* mutant murine CMT167 cells expressing doxycycline-inducible control shRNA or two independent shRNAs targeting *Urod.* (**B**) Schematic illustration of CMT167 syngeneic experiment. (**C**) Quantification of tumor volumes of CMT167 cells expressing the indicated shRNA sequence transplanted in C57BL/6J mice (n=15 mice per group, *scrambled* shRNA data are shown in **Supplementary Fig. 4**). Graph represents mean tumor volumes and error bars represent standard error of the mean. (**D**) Uniform manifold approximation and projection (UMAP) analysis of tumor infiltrating lymphocytes (TILs) colored by cell types. (**E**) Quantification of TILs (expressed as a % of CD45^+^ cells) from flow mass cytometry (mass CyTOF). n=5 *scrambled* mice, n=5 *Urod* shRNA mice, n=5 *Urod* shRNA + ISRIB, n=4 *Urod* shRNA + αPD-1, n=5 *Urod* shRNA + ISRIB +αPD-1. Graphs represent mean values and error bars represent standard deviation from the mean. A Student’s t-test was used to determine statistical significance (* p<0.05, ** p<0.005, *** p<0.0005).

We next investigated the immune microenvironment of these tumors utilizing mass CyTOF flow cytometry. Cells were analyzed with traditional gating strategies (**Supplementary Fig. S4D-F**), and immune populations were identified as Cisplatin-, CD45^+^ (live CD45^+^ cells), and further gated by cell type (**Supplementary Fig. S4D-E**) as a percentage of live CD45^+^ cells. Uniform manifold approximation and projection (UMAP) analysis of tumor infiltrating lymphocytes (TILs) was obtained and annotated by cell type (**Supplementary Fig. S5A**). The most significant changes were observed between *Urod* shRNA tumors and *Urod* shRNA tumors with ISRIB and PD-1 blockade (**Fig. 5D**). Analysis of individual immune subsets revealed a three-fold decrease in CD8^+^ T cells between *scrambled* and *Urod* shRNA tumors and a four-fold decrease in Granzyme B^+^ CD8^+^ T cells (**Fig. 5E**), consistent with our *in vitro* data identifying CD8^+^ T cells as a key immune subset affected by ISR activation in tumor cells. As expected, PD-1 blockade improved the infiltration of CD8^+^ T cells and Granzyme B^+^ CD8^+^ T cells in *Urod* shRNA tumors but did not have significant effects on any other immune subsets. ISRIB treatment alone did not significantly impact immune cell populations, however the combination of ISRIB and PD-1 blockade exhibited the most significant effects when considering all cell populations. The combined therapies did not further increase the infiltration of CD8^+^ T cells and Granzyme B^+^ CD8^+^ T cells compared with PD-1 blockade alone, but the combination did ehance dendritic cell (DC) infiltration (**Fig. 5E**). Moreover, the combination significantly reduced regulatory T cells (T_reg_), myeloid derived suppressor cell (MDSC), and natural killer (NK) cell populations compared to untreated tumors. No significant changes were observed for total CD3^+^ T cell populations, CD4^+^ T cells, B cells or tumor-associated macrophages (**Supplementary Fig. S5B**). Overall, the combined treatment of ISRIB and PD-1 blockade enhanced CD8^+^ T infiltration and cytotoxic activation to a similar extent as PD-1 blockade alone, and significantly enchanced antigen presentation through DC enrichment, while suppressing traditionally immunosuppressive immune subsets including T_reg_ and MDSC populations. These data suggest that the combination of ISRIB and PD-1 blockade may improve therapeutic responses of NSCLC patients with tumors expressing CD155 and PD-L1.

### Correlated expression of CD155 and PD-L1 in primary human lung adenocarcinomas

We next evaluated the correlation of CD155 and PD-L1 in two independent patient cohorts. In a panel of 33 primary human lung adenocarcinomas and matched normal tissue provided by the UT Southwestern Tissue Management Shared Resource (TMSR), H-score analysis of CD155 and PD-L1 revealed a significant positive correlation (Pearson correlation r = 0.506, p = 0.0031) with co-expression of both checkpoints observed in approximately 30% of tumors (**Fig. 6A-B**).

**Figure 6:**
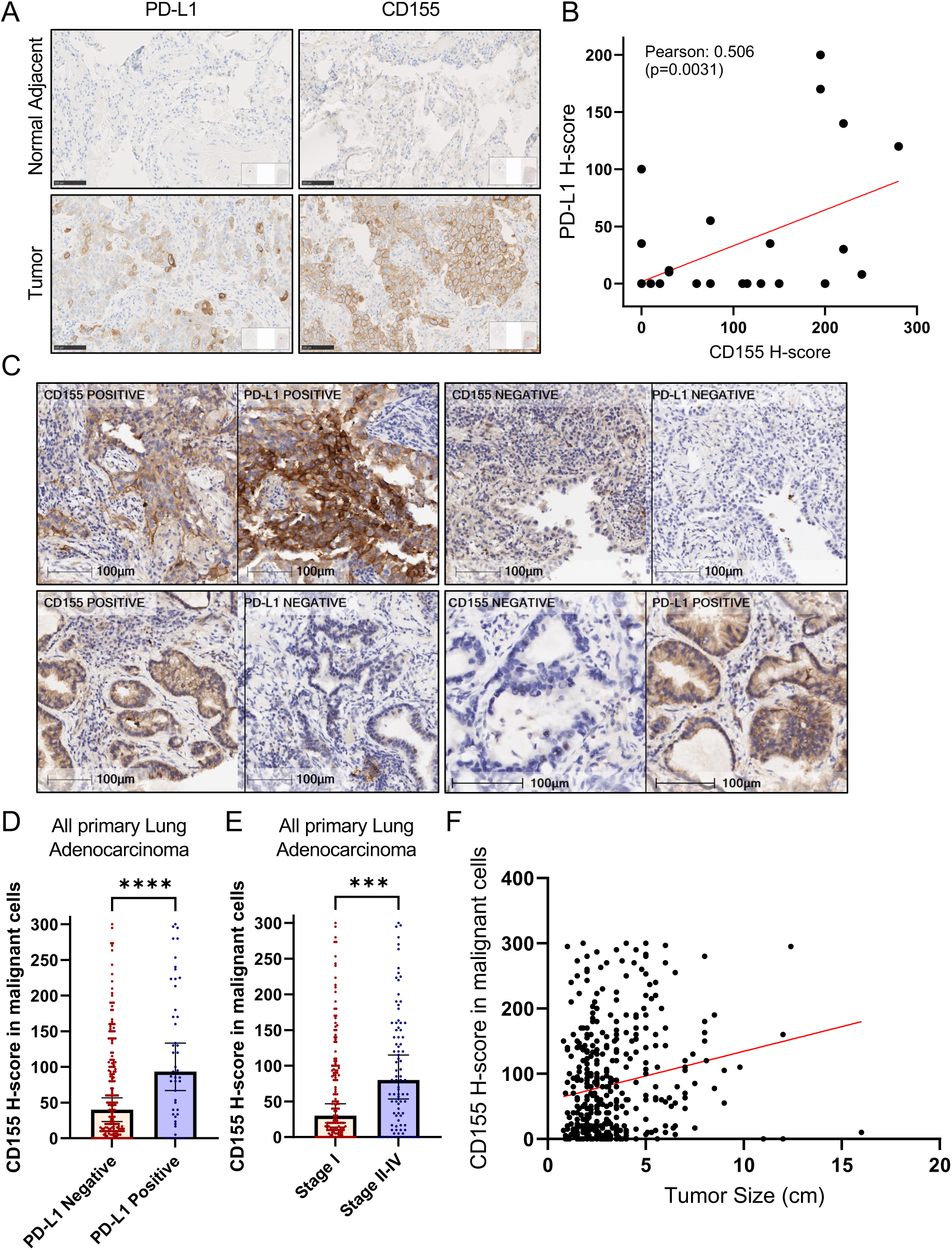
CD155 and PD-L1 are positively correlated in primary human lung adenocarcinoma tumors. **(A)** Representative images of IHC for PD-L1 and CD155 of 33 primary human lung adenocarcinomas. 20X representative images shown, scale bar=100μm (**B**) Correlation of CD155 H-score and PD-L1 H-score of tissues in (**A**) (Pearson correlation r = 0.506, p = 0.0031). (**C**) Microphotographs of CD155 and PD-L1 immunohistochemistry analysis of primary lung adenocarcinomas displaying different combinations of expression. (CD155+/PD- L1 +, CD155- /PD-L1 -, CD155+ /PD-L1-, CD155-/PD-L1 +). 20X representative images shown, scale bar=100μm. (**D**) Association of CD155 H-score expression in all primary lung adenocarcinoma with PD-L1 status (cut-off for positive PD-L1 expression: Tumor proportion score >=1%) (p= 0.0004). A Mann Whitney test was used to determine statistical significance. Graph represents the median; error bars represent the 95% confidence interval. (**E**) Association of CD155 H-score expression in all primary lung adenocarcinoma with pathological stage (p= 0.001). A Mann Whitney test was used to determine statistical significance. Graph represents the median; error bars represent the 95% confidence interval. (**F**) Correlation of CD155 H-score expression with tumor size (Spearman correlation r= 0.1946, p<0.0001).

Next, we performed immunostaining with a CD155 antibody on a panel of over 400 clinically and pathologically annotated primary NSCLCs utilizing a tissue microarray (TMA) (**Supplementary Table S1**). We observed similar trends in CD155 and PD-L1 staining (**Fig. 6C, Supplementary Fig. S6A**), and categorized tumors as PD-L1 positive or negative to perform analysis with CD155 H-score. CD155 H-score was significantly associated with PD-L1 positivity in primary lung adenocarcinomas (**Fig. 6C-D**) and all primary NSCLC (**Supplementary Fig. S6B**), but not in squamous cell carcinoma (data not shown). CD155 expression was further elevated in squamous cell carcinoma as compared to adenocarcinoma (**Supplementary Fig. S6C**), in males versus females (**Supplementary Fig. S6D)**, and was associated with smoking status (**Supplementary Fig. S6E**). Additionally, CD155 expression was significantly associated with higher tumor stage (**Fig. 6E, Supplementary Fig. S6F)**, and tumor size (**Fig. 6F**). In summary, our data support a model (**Fig. 7**) wherein stresses present in the tumor microenvironment activate the ISR pathway in tumor cells. This results in increased translation of two key immune checkpoint proteins, CD155 and PD-L1, which suppresses T cell function by binding the inhibitory T cell receptors, PD-1 and TIGIT respectively, leading to enhanced tumorigenesis. Thus, the combination of ISR inhibition with PD-1 blockade may represent a new treatment modality to improve current strategies.

**Figure 7:**
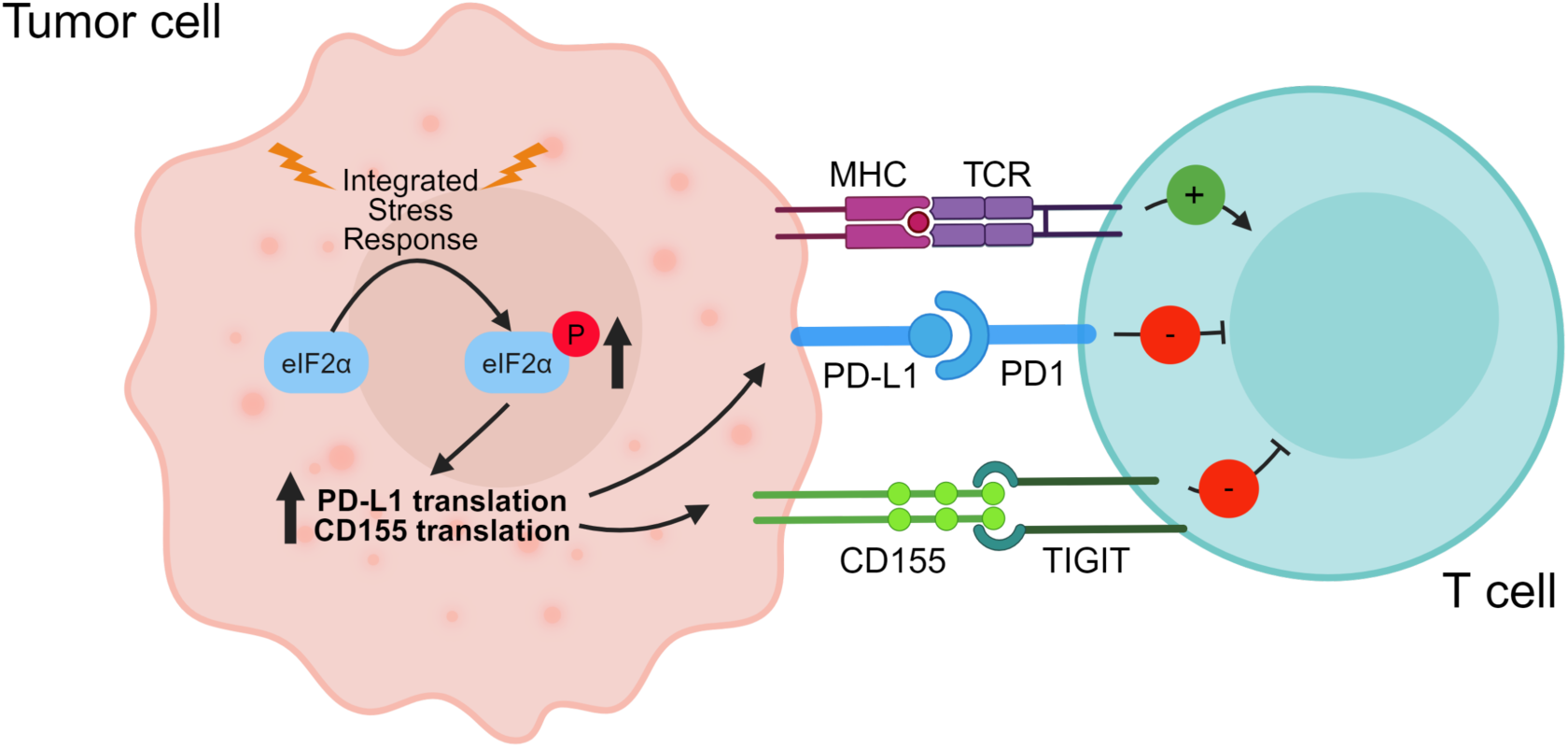
Model of translational control of CD155 and PD-L1 in response to ISR activation. Stresses commonly present in the tumor microenvironment activate the ISR pathway in tumor cells. This results in enhanced translation of *CD155* and *PD-L1*, which suppresses T cell function by binding the inhibitory T cell receptors PD-1 and TIGIT respectively, thereby leading to enhanced tumorigenesis. Figure created with Biorender.com.

## Discussion

Immune checkpoint blockade is currently a first line therapy for lung cancer patients, yet only ∼30% of patients have durable responses. A thorough understanding of both intrinsic and extrinsic mechanisms of immune checkpoint regulation is essential for improving patient responses. In this study, we found that activation of the ISR pathway enhances translation of CD155 and PD-L1 through the bypass of inhibitory uORFs in their 5′ UTR, leading to coordinated expression of both immune checkpoint proteins in human and mouse lung cancer cells. This results in inhibition of T cell activation and cytotoxic activity *in vitro* and *in vivo*. Furthermore, we found that the efficacy of PD-1 blockade is enhanced with the ISR inhibitor ISRIB. Importantly, the combination of ISRIB with anti-PD-1 significantly reduced the percentage of immunosuppressive immune cell subsets compared with PD-1 blockade alone, while enhancing antigen presenting and anti tumor immune cell populations. This combinatorial approach may represent a new treatment strategy, particularly for patients with tumors that co-express CD155 and PD-L1. Interestingly, several recent studies have shown that CD155 expression is associated with resistance to anti-PD-1 therapy in NSCLC (39) and melanoma patients (40). These observations are consistent with our results demonstrating that expression of CD155 and PD-L1 frequently co-occur in lung cancer. Thus, our results provide new insight into mechanisms of tumor immune escape.

The ISR represents a targetable pathway that may improve immunotherapies. ISRIB was first identified as memory-enhancing drug with limited toxicity (33), and restores translation by binding and enhancing eIF2B activity. ISRIB has demonstrated anti-tumor efficacy in multiple cancer models. For example, in a PDX model of advanced castration-resistant prostate cancer, ISRIB treatment reduced metastatic progression and selectively triggered cancer cell cytotoxicity (41). The ISR has also been shown to modulate breast cancer cell plasticity through 5′ UTR regulation of cell fate and lineage markers *NANOG, SNAIL* and *NODAL*. ISRIB treatment reversed this cancer stem cell switch and bolstered chemotherapy in a triple-negative breast cancer PDX model (42). ISRIB diminishes anchorage independent growth and migration in lung adenocarcinoma cell lines (43), and inhibits tumorigenesis in both subcutaneous and orthotopic *Kras^G12C^* lung tumor models (15). While these prior studies did not focus on the effects of ISRIB on the immune system, our work demonstrates that inhibiting the ISR pathway also enhances anti-tumor immunity, especially in combination with PD-1 blockade. Given that ISRIB derivatives are currently in clinical trials for cognitive disorders, these compounds are poised to enter clinical trials for the treatment of non-small cell lung cancers.

Targeting of the CD155/TIGIT axis has gained attention as a promising adjuvant to PD- L1/PD-1 blockade with several groups demonstrating that the CD155/TIGIT axis is linked to immune evasion, including in pancreatic ductal adenocarcinoma (PDAC) (28, 39, 40, 44, 45). Interestingly, the neoantigen-specific immune response in PDAC is linked to this immune axis, and the combination of TIGIT/PD-1 blockade with a CD40 agonist antibody (which aids the recruitment of immune cells) resulted in increased CD8^+^ T cell infiltration and reduced MDSC populations, thus eliciting anti-tumor responses in the traditionally “cold” PDAC tumor microenvironment (27). This and other studies provided proof-of-principle for the development of monoclonal antibodies targeting TIGIT (46) with multiple human anti-TIGIT monoclonal antibodies currently in clinical trials in combination with anti-PD-1/PD-L1 antibodies. For example, Genetech’s tiragolumab (anti-TIGIT antibody) plus atezolizumab (anti-PD-L1) trial (CITYSCAPE) is ongoing. Initial results demonstrated that tiragolumab plus atezolizumab showed an improvement in objective response rate (ORR) and progression-free survival (PFS) in recurrent or metastatic PD-L1-positive NSCLC compared with placebo plus atezolizumab, with the most significant responses noted in patients with high PD-L1 expression (47). Analysis of patient serum from the CITYSCAPE trial found that responsive patients exhibited an “inflamed” tumor myeloid phenotype. In mouse models, the tiragloumab antibody enriched the same cell types through FcyR interactions, driving exhausted CD8+ T cells to transition to a more memory-like T cell phenotype (44). Another ongoing phase 2 trial, ARC-7, similarly demonstrated enhanced anti-tumor activity of combined zimberelimab (anti-PD-1) with domvanalimab (an Fc-silent anti-TIGIT antibody) in metastatic PD-L1-high NSCLC, in comparison to PD-1 blockade alone (48). Despite these promising results, several other recent trials did not meet their primary endpoints (49, 50). It is important to note that the biological differences in the patient populations enrolled in these trials may contribute to mixed results, and further trials are ongoing or in recruitment phases. These studies highlight the importance of future studies to clarify mechanisms of immune checkpoint regulation and immune escape within lung cancer subtypes.

Here we uncover a novel mechanism of immune checkpoint activation and illuminate the role of translation in the regulation of the CD155 and PD-L1 immune checkpoint proteins. Our findings suggest that ISR inhibition may enhance the combined PD-1/TIGIT blockade. Further investigation of the extent to which targeting the ISR pathway synergizes with immune checkpoint blockade and overcomes immunotherapy resistance will be an important priority for future work.

## Materials and Methods

### Cell culture

All human lung cancer cell lines (obtained from Dr. John Minna) were cultured in RPMI 1640 media (Gibco, A1049101) supplemented with 5% FBS (Sigma, F2442) and 1% antibiotic-antimycotic (anti-anti, Invitrogen, 15240-062). WT and eIF2α mutant Mouse Embryonic Fibroblasts (MEFs), from Dr. Randal Kaufman, were cultured in DMEM (Gibco, 11995073) supplemented with 10% FBS (Sigma, F2442), 1% anti-anti, 2mM l-glutamine (Thermo Fisher, 25030081), 2% MEM amino acid solutions (Gibco, 11140050) and 1mM Sodium pyruvate (Sigma, P2256-100G). CMT167 and Jurkat T cells were cultured in RPMI 1640 media supplemented with 10% FBS and 1% antibiotic-antimycotic (anti-anti, Invitrogen, 15240-062). JAWS II (obtained from ATCC) cells were cultured in IMDM (Gibco, 12440053) with 10% FBS, 4 mM l-glutamine, 1% antibiotic-antimycotic, 0.5 mM 2-ME, 1 mM sodium pyruvate, and 5 ng/ml murine GM-CSF (Abcam, ab259385). Primary mouse bone marrow dendritic cells were isolated from the femurs of C57BL/6J mice and grown in RPMI 1640 media (Gibco, A1049101) supplemented with 10% FBS (Sigma, F2442), 20 ng/ml murine GM-CSF, and 1% antibiotic-antimycotic (anti-anti, Invitrogen, 15240-062). Human PBMCs (obtained from AllCells) were cultured in RPMI 1640 media supplemented with 10% FBS and 1% antibiotic-antimycotic (anti-anti, Invitrogen, 15240-062). For ISR pathway activation and inhibition, cells were treated with 100uM or 200uM Salubrinal (Tocris, 23-471-0) and/or 500-800nM ISRIB (Fisher Scientific, 5284) for 24h or 48h. For ISR activation, LUAD cells were treated with thapsigargin (5uM, Sigma, T9033), or with Arsenite (10-50uM), or grown in a hypoxic chamber for 24 or 48h. For amino acid deprivation, cells were grown in RPMI 1640 media without amino acids (US Biological, R9010-01) supplemented with 5% FBS (Sigma, F2442) and 1% antibiotic-antimycotic (anti-anti, Invitrogen, 15240-062). All chemicals are listed in **Supplementary Table 2**. All cell lines were tested and found to be mycoplasma free using a direct PCR method with GoTaq Green Master Mix (Promega, M712).

### Immune cell co-cultures and ELISA assays

H358 cells (5×10^4^) were plated into 24-well plates 24 hours before co-culture. The next day media was removed, and Jurkat T cells (4×10^5^) or primary human PBMCs (1×10^5^) were added in fresh media to H358 cells. After 1 hour, CD3 (Thermo Fisher, 16-0037-85, RRID:AB_468855) and CD28 (Thermo Fisher, 16-0289-85, RRID:AB_468927) (4ug/ml each for Jurkat T cells, 1ug/ml each for PBMCs) activating antibodies were added and cells were co-cultured for an additional 24 hours. Conditioned media was then collected, centrifuged to remove cells and debris, and immediately used for IL-2 (Abcam, ab100566) and Granzyme-B ELISA assays (Abcam, ab235635) according to manufacturer’s instructions.

### Plasmids

LentiCRISPR V2 (#52961, RRID:Addgene_52961), PAX2 (#12260, RRID:Addgene_12260) and MD2 (#12259, RRID:Addgene_12259) plasmids were obtained from Addgene. The pTRIPZ plasmid was obtained from Dharmacon (RHS6371, RRID:Addgene_206981). Generation of knockout cell lines using CRISPR-Cas9.: HEK 293T cells (RRID:CVCL_0063) were seeded in 15 cm dishes and co-transfected with lentiCRISPR V2 (1ug) and PAX2 (6.6ngg), MD2 (3.3ng) helper plasmids using Lipofectamine 3000 (Invitrogen, L3000150). Lentiviral supernatant was collected 48h post transfection, filtered, and concentrated with Lenti-X Concentrator (Takara, 631232). Recipient cells were infected overnight with 1/10 of viral concentrate in media containing 8ug/mL polybrene (Tocris, 7711/10). 24-48h later, transduced cells were cultured in fresh media containing 1ug/mL puromycin (Gibco, A1113803) for 10-12 days.

### RNA extraction and qRT-PCR analysis

Total RNA was isolated from cells using the RNeasy Mini Kit (Qiagen, 74106) according to manufacturer’s instructions. For qRT-PCR of mRNA, cDNA synthesis was performed with 1-2μg RNA for reverse transcription using Superscript IV Vilo Master Mix (5X) (Invitrogen, 11756500). mRNA expression was assessed using quantitative real-time PCR with 2X SYBR Green Fast qPCR Mix (Abclonal, RK21203). mRNA levels were normalized to β-actin or luciferase expression, with gene expression levels measured using the ΔΔCT method. PCR primers, designed to cover exon-exon junctions, are provided in **Supplementary Table 3**. To monitor mRNA decay, cells were treated with Actinomycin D (10ug/ml) (Thermo Fisher, A7592) to halt transcription and RNA was isolated.

### Western blotting

Cells and tissues were lysed in RIPA (Invitrogen, 89901) buffer containing Halt Protease/Phosphatase Inhibitor cocktail (Fisher Scientific, PI78442) and homogenized using a Bioruptor (Diagenode). Proteins were quantified using the Bicinchoninic Acid (BCA) assay (Thermo Fisher, 23225), subject to separation using NuPage Bis-Tris gels (Invitrogen, NW04120BOX) and transferred to a nitrocellulose membrane. The membranes were blocked for 1h at RT in 5% milk and probed with primary antibodies in 5% milk overnight at 4°C. After incubating the membrane with the appropriate secondary antibody conjugated to horseradish peroxidase, protein levels were detected with SuperSignal Extended Dura substrate (Thermo Scientific, 34076). Antibodies are listed in **Supplementary Table 4**. To monitor protein degradation, cells were treated with Cycloheximide (40ug/ml, Sigma, C7698), cells were lysed in RIPA buffer, and protein degradation was measured through western blot. Western blots were quantified using ImageJ analysis (RRID:SCR_003070) and normalized to loading controls (Vinculin or GAPDH).

### Flow Cytometry

Confluent cells were trypsinized and centrifuged at 300g for 5 minutes. Cells were counted (Countess Automated Cell Counter) and washed with cell staining buffer (BioLegend, 420201). Cells were incubated in the dark for 20 minutes on ice in cell staining buffer containing APC-PD- L1 antibody (BioLegend, 329708) or FITC-CD155 antibody (Biolegend, 337628) at a concentration of 0.4ug antibody/million cells. Stained cells were centrifuged at 300g for 5m and washed with cell staining buffer twice. Final cell pellets were resuspended at a cell density of 2×10^6^ cells/ml in 3% FBS in PBS and analyzed at the UTSW Flow Cytometry Core Facility (FACSCalibur, BD Biosciences). FlowJo software was used to analyze mean fluorescent intensity (RRID:SCR_008520).

### Transient knockdown using siRNA

Recipient cells were seeded in 6 well plates. The next day, cells were transfected with siRNA pools (5uM, siGENOME Dharmacon pools targeting UROD (M-013415-01-0005), or non-targeting control (D-001206-13-20)) and Dharmafect solution 4 (T-2004-02) in serum-free media according to manufacturer’s instructions. Cells were replenished with fresh complete media the next day and harvested 48-72h later for RNA/western analysis.

### Inducible knockdown of *Urod*

HEK 293T cells were co-transfected with pTRIPZ (Dharmacon, RHS5087) with helper plasmids as described above. CMT167 cells were infected overnight with *scrambled* or *Urod* shRNA (2 independent shRNAs) concentrated virus and 8ug/mL polybrene (Sigma). Transduced cells were selected in 2ug/mL puromycin (Gibco, A1113803) or 4ug/mL blasticidin (Thermo Fisher, R21001) for 1 week and cultured in 2-3ug/mL doxycycline (RPI, D43020-250.0) for 1-2 days. Cells were harvested for RNA/protein to assess knockdown.

### Tumorigenesis assays

CMT167 cells (3×10^5^) expressing a *scrambled* shRNA or *Urod* shRNA, or non-transfected CMT167 cells were injected subcutaneously into the right flanks of 6-8-week-old C57BL/6J female mice (Jackson laboratory, RRID:IMSR_JAX:000664). For Salubrinal treatment, mice were injected with 1mg/kg (daily, IP) or vehicle (0.2% v/v Tween80 in 1X PBS), once tumors were palpable. For ISRIB treatment, mice were injected with 2mg/kg (daily, IP) or vehicle (0.5% w/v hydroxypropylmethylcellulose dissolved in 1X PBS plus 0.2% v/v Tween80, adjusted to pH 4), once tumors were palpable. For PD-1 blockade, beginning on Day 7 mice were injected with 200ug antibody IP every three days. For tumorigenic studies with CMT167 cells expressing a *scrambled* shRNA or *Urod* shRNA, mice were kept on doxycycline water (2g/L doxycycline (RPI, D43020-250.0), 2% sucrose, (Fisher Scientific, 50-188-2396)) beginning on Day 7 and continuing for the duration of the experiment. Tumor volume was measured using calipers every 3 days. Tumor volume was calculated using the formula (length x width^2^)/2. Mice were randomized after injection of tumor cells, and prior to initiation of drug or antibody treatments.

### Mass CyTOF flow cytometry analysis of tumor infiltrating lymphocytes

Tumors were excised from euthanized mice and homogenized using a Tissue Dissociator. Tumor cells were digested at 37°C at 150 rpm for 30 minutes in digest buffer (RPMI 1640 containing 10% FBS, 1 mg/mL Collagenase A (Sigma, 10103586001) and 5 ug/mL DNAse I(Sigma, 10104159001)). All subsequent steps were performed on ice. Digested tumors were filtered using a 70um cell strainer, pelleted at 500xg, and incubated with Red Blood Cell lysis buffer (Sigma, 11814389001). Cells were washed twice, and then processed for Helios Mass CyTOF (Fluidigm) according to manufacturer’s instructions. Antibodies are listed in **Supplementary Table 4**. Briefly, cells were stained with Cisplatin (5uM) for 5 minutes at room temperature, quenched with Maxpar Cell staining buffer (Fluidigm, 201068), washed and counted. 1-3 million cells were analyzed per sample. The cells were centrifuged and resuspended in 50uL Fc Receptor Blocking Solution, incubated for 10 minutes at room temperature, followed by the cell surface antibody cocktail for 30 minutes at room temperature. Cells were washed and centrifuged twice, fixed with 1X Maxpar Fix I Buffer (Fluidigm, 201065) for 30 minutes at room temperature, washed twice and resuspended and incubated in the cytoplasmic/secreted antibody cocktail for 30 minutes. Cells were washed and centrifuged twice, then incubated with Invitrogen FOXP3 Fixation/Permeabilization solution for 30 minutes, followed by two more washes and incubation with the nuclear antigen antibody cocktail for 30- 45 minutes. Cells were then washed and fixed with a fresh 1.6% formaldehyde solution for 10 minutes and then centrifuged and stained overnight with 125nM Cell-ID Intercalator-Ir at 4°C. The next day, cells were washed twice with Maxpar Cell Acquisition Solution (Fluidigm, 201240), filtered through a nylon membrane into flow cytometry tubes for analysis, and pelleted. Samples were analyzed at the UTSW Flow Cytometry Core Facility (Helios, Standard BioTools). FlowJo software was used to gate live, single cells. OMIQ (omiq.ai) was then used to calculate immune cell populations (%) of CD45+ live cells.

### Multiplex fluorescent Immunohistochemistry (IHC-F)

Paraffinized tissues were dewaxed with xylene (3 x 10 minutes) and rehydrated with a sequential graded series of ethanol solutions (100%, 95%, 70%, 10 minutes each). After rehydration, slides were rinsed in DI water and fixed in 10% neutral buffered formalin for 20 minutes. Slides were then processed for multiplexed fluorescent IHC according to manufacturer’s instructions (Opal 4-Color Manual IHC Kit, Akoya Biosciences, NEL810001KT). Briefly, slides were microwaved for 30 minutes in Antigen Retrieval buffer, blocked in blocking solution for 10 minutes, then incubated in primary antibody dilutions overnight at 4°C. The next day, the slides were washed and incubated with Secondary-HRP, washed again, and incubated with Opal fluorophore working solution for 10 minutes. This process was then repeated beginning with microwaving in Antigen Retrieval buffer until all targets were detected. Finally, slides were counterstained with DAPI, and coverslips mounted (Prolong Diamond, Invitrogen, P36961) and cured overnight before imaging. Slides were scanned at the UTSW Whole Brain Microscopy Facility (WBMF) on Zeiss Axioscan 7. IHC-F images were quantified using ImageJ analysis (RRID:SCR_003070).

### Immunohistochemistry (IHC) of tumor tissues

#### UT Southwestern

Immunohistochemical analysis was performed on a Dako Autostainer Link 48 system. Briefly, the slides were baked for 20 minutes at 60°C, then deparaffinized and hydrated before the antigen retrieval step. Heat-induced antigen retrieval was performed at pH 9 for 20 minutes in a Dako PT Link. The tissue was incubated with a peroxidase block and then with the antibodies EIF2S1 (1:150 for 20 minutes, Abcam, ab32157, RRID:AB_732117), CD155 (1:100 for 20 minutes, CST, 81254S, RRID:AB_2799970), PD-L1 (1:100 for 30 minutes, CST, 13684S, RRID:AB_2687655). The staining was visualized using the EnVision FLEX visualization system. Slides were further scanned for analysis on a Hamamatsu Nanozoomer 2.0HT. H-score analysis was performed by a trained pathologist, Dr. Bret Evers.

#### MD Anderson tissue microarray

A tissue microarray (TMA) with 410 surgical resected primary non-small cell lung carcinomas (NSCLC) collected from 2006 to 2009 at The University of Texas MD Anderson Cancer Center (MD Anderson; Houston, TX, USA) was used in this study. The TMA was constructed using three 1-mm width cores per each sample. This study was approved by the MD Anderson Institutional Review Board and was conducted according to the principles of the Helsinki Declaration. Annotated clinicopathological information, including demographics, smoking history, histological diagnosis, and pathologic tumor-node-metastasis stage (TNM staging system) was available for all samples, and summarized in **Supplementary Table 1**.

#### Immunohistochemistry staining

Immunohistochemistry staining for CD155 (CST, 81254, RRID:AB_2799970) was performed using an automated staining system. The immunohistochemistry protocol is briefly described as follows: tissue sections (4 μm) were stained in a Leica BOND RX automated stainer (Leica Biosystems Nussloch GmbH). The tissue sections were deparaffinized and rehydrated following the Leica BOND protocol. Antigen retrieval was performed for 20 min with Bond Epitope Retrieval Solution #2 (Leica Biosystems, equivalent EDTA, pH 9.0). Primary antibody (dilution at 1: 100) was incubated for 15 min at room temperature and detected using the Bond Polymer Refine Detection kit (Leica Biosystems, Cat# DS9800) with DAB as chromogen. The slides were counterstained with hematoxylin, dehydrated, and cover slipped. PD-L1 (E1L3N) antibody immunohistochemistry procedure was previously reported, and slides were available for immunohistochemistry evaluation (48).

### Immunohistochemistry assessment of CD155 and PD-L1

Immunohistochemistry-stained slides were scanned using the Aperio AT2 scanner (Leica biosystem) and visualized using Aperio Image Scope software (v 12.3.3.7026). CD155 immunostaining was evaluated in the membrane of malignant cells (MCs) by two independent pathologist (MS and LS) and quantified using a 4-value intensity score [0+ (no staining), 1+ (weak staining), 2+ (moderate staining), and 3+ (strong staining)] and the percentage (0%–100%) of the extent of reactivity. Membrane PD-L1 was evaluated by two pathologists (MS and LS) using standard microscopy and reported as percentage of MCs with positive expression. Positive PD-L1 expression was defined using a cutoff of ≥1% of staining in MCs.

### Polysome Profiling

Sucrose gradients (5% to 50%) were prepared in advance with the BioComp Gradient Master and stored at 4°C overnight. The next day, 20-40×10^6^ cells/sample were treated for 5 minutes with 100ug/ml cycloheximide and trypsinized. Cells were then washed twice with ice-cold PBS (second wash containing 100ug/mL cycloheximide). Cells were pelleted at 200xg for 5 min at 4°C and resuspended in 500ul of Polysome Extraction Buffer (5mM Tris-HCl (pH 7.5), 1.5mM KCl, 2.5mM MgCl2, 0.5% Triton X-100, 0.5% Sodium Deoxycholate, 2mM DTT in distilled water) containing cycloheximide, protease inhibitor cocktail (Fisher Scientific, PI87785) and RNAse inhibitors (Promega, N2515). Cells were vortexed briefly, incubated on ice for 10 minutes, sheared through a 27.5 gauge needle three times, and lysates were centrifuged at 15,000 rpm for 5 min at 4°C and the supernatant lysate RNA concentration quantified by Nanodrop. An equal amount of lysate (500-600ug RNA) was loaded across all gradients. The gradients were centrifuged at 35,000 rpm for 2h at 4°C and run on a fractionator machine (BioComp Piston Gradient Fractionator) to visualize and collect polysome fractions. Each collected fraction was mixed with 3x volume of 100% ethanol and 20ug glycogen carrier (Roche, 10901393001) and incubated overnight in −20°C. The next day, fractions were centrifuged at 20,000g for 30m at 4°C to precipitate RNA pellets. Pellets were dried for 20 min at RT, resuspended in 100uL Nanopure water and 350uL RNeasy RLT lysis buffer and loaded onto RNeasy columns. The RNeasy kit was used to isolate RNA and cDNA synthesis and real-time PCR performed. 20ng of Luciferase mRNA control (Promega, L4561) was added to each fraction prior to RNA extraction to control for variability in total RNA in fractions during RNA isolation and reverse transcription. Fractions associated with <3 ribosomes were grouped together (poorly translated mRNAs) and fractions with >3 ribosomes were grouped (efficiently translated mRNAs).

### Dual luciferase assays

50×10^3^ MEF cells were seeded/well in 12-well plates in triplicate and transfected with a Renilla plasmid (20ng, Promega, E2241), Firefly luciferase pGL3 plasmid (Promega, E1751, RRID:Addgene_212936) expressing CD155 wildtype 5′ UTR or various mutant constructs (200ng) and a carrier pUC19 plasmid (400ng) per well using Fugene HD (Promega, E2311) at a 3:1 Fugene:DNA ratio. Luciferase activity was measured 48h post transfection using a Luminescence plate reader (Promega, E1910). Firefly luciferase activity was normalized to Renilla luciferase activity to obtain Relative luciferase levels/sample.

### Statistics and reproducibility

A Student t-test was used for comparisons between two groups with normal data distribution (for real time qPCR and other indicated analyses) or with Welch’s correction (for comparing % CD8+ T-cells and other immune subsets). Chi-square or Fisher’s exact tests were conducted to investigate if there were significant differences in clinicopathological characteristics between the CD155 membrane IHC negative and positive groups. For tumor implantation assays, we used generalized linear mixed models (R package nlme version 3.1.140) to examine if there were significant differences in tumor volume over time among the treatment groups. Reported p-values were adjusted for multiple comparisons using Bonferroni corrections. Representative results from at least two independent experimental repeats are shown, except where specified otherwise in the figure legends.

## Ethics statement

All procedures involving mice were performed in accordance with the recommendations of the Panel on Euthanasia of the American Veterinary Medical Association and protocols approved by the UTSW Institutional Animal Care and Use Committee. Mice were monitored closely throughout all experimental protocols to minimize discomfort, distress or pain. C57BL/6J mice were obtained from The Jackson Laboratory.

## Data Availability

Raw data for **Fig. 6C-F** and **Supplementary Fig. 6** were generated at UT MD Anderson Cancer Center, in the Department of Translational Molecular Pathology. All primary data supporting the findings of this study are available from the corresponding author upon request.

## Acknowledgements

We thank Dr. Randal Kaufman (SBP Discovery Institute) for sharing eIF2α wild-type (S/S) and mutant (A/A) MEFs, members of the Mendell laboratory for assistance with polysome profiling, and Angie Mobley for assistance with flow cytometry. We thank Joshua Mendell and members of the O’Donnell laboratory for critical reading of the manuscript. K.A.O. is supported by the NCI (R01CA273585, R01CA207763, and P50CA70907), the Cancer Prevention Research Institute of Texas (CPRIT RP190610 and RP200327), the Welch Foundation (I-1881), the V Foundation (T2021-011), and the Department of Defense (DoD LC190249). S.T.J. is supported by the NCI (F32CA274982). SS is supported by a CSIR RDSF grant (IHP24002) and a Melanoma Research Foundation Career Development Award. Slide scanning was made possible on Zeiss Axioscan 7 courtesy of the following funding (1S10OD032267-01, to Denise Ramirez) and the Whole Brain Imaging Core at UTSW. We also acknowledge the assistance of the UTSW Tissue Management Shared Resource, a shared resource at the Simmons Comprehensive Cancer Center, which is supported in part by the National Cancer Institute P30 CA142543.

## Supplementary Figure Legends

**Supplementary Figure 1: The effect of ISR pathway activation on multiple immune checkpoint proteins.** (**A**) Overview of cell surface immune checkpoint proteins and their corresponding receptors on T cells. PD-L1 and PD-L2 (Programmed death ligand 1 and 2) are ligands for PD-1 (Programmed cell death protein 1) whose binding inhibits both TCR and CD28 signaling on T cells, suppressing T cell function. CD155 (Cluster of differentiation 155) engages with several immune cell receptors to facilitate immune suppression, including TIGIT (T Cell Immunoreceptor with Ig and ITIM Domains), which is expressed on T cells, dendritic cells, and NK cells. Galectin-3 is a secreted member of the galectin family capable of binding both LAG3 (Lymphocyte-activation gene 3) and CTLA-4 (Cytotoxic T-lymphocyte associated protein 4) to inhibit T cell activity. Galectin-9 binds TIM-3 (T-cell immunoglobulin and mucin-domain containing-3) induceing T-cell apoptosis. HVEM (Herpes virus entry mediator) binds BTLA (B and T lymphocyte attenuator) to suppress T cell function or bind LIGHT (tumor necrosis factor superfamily member 14, not shown) to induce activation depending on the microenvironment. (**B**) Western blot analysis of human H441, H358, PC9, and HCC827 cells treated for 24 hours with 5uM thapsigargin (ER Ca+ ATPase pump inhibitor) to induce ER stress, or DMSO vehicle control. (**C**) Quantitative real-time PCR analysis of *PD-L1(CD274), CD155(PVR), GADD34*, and *UROD* in H358 cells transfected with control or *UROD* sgRNA. (**D**) Quantitative real-time PCR analysis of *PD-L1(CD274), CD155(PVR),* and *GADD34,* in H358 cells treated for 24 hours with 100uM Salubrinal or DMSO vehicle control or (**E**) grown for 24 hours in RPMI 1640 media supplemented with or without amino acids. Target transcript mRNA was normalized to *Actin*, and fold change was calculated using the ΔΔCT method. Error bars represent standard deviation from the mean from n=3 biological replicates. Data from a single experiment are shown and representative of two independent experiments. A Student’s t-test was used to determine statistical significance (* p<0.05, ** p<0.005).

**Supplementary Figure 2: The effect of ISR pathway activation on protein and mRNA stability, and translation**. (**A**) Quantitative real-time PCR analysis of *PD-L1(CD274)* in individual ribosomal fractions from (**Main Figure 2A**). (**B**) Quantitative real-time PCR analysis of *ATF4* in pooled (<3 or >3 ribosomes) and individual (**C**) ribosomal fractions using a single primer pair set covering an exon-exon junction. (**D**) Quantitative real-time PCR analysis of *CD155(PVR)* in individual ribosomal fractions from (**Main Figure 2A**). (**E**) Quantitative real-time PCR analysis of *CD155(PVR)* in pooled (<3 or >3 ribosomes) and individual (**F**) ribosomal fractions from a second primer set (Primer 2) covering a different exon-exon junction. *PD-L1, ATF4,* and *CD155* mRNA expression in each fraction was normalized to *Luciferase* and mRNA abundance was calculated as the % of total in all fractions. Luciferase control mRNA was added to each fraction prior to RNA extraction to control for variability. Error bars represent standard deviation from the mean from three technical replicates of three independent fractions (<3 or >3 ribosomes). Fractions associated with <3 ribosomes were grouped to represent poorly translated mRNAs, fractions associated with >3 ribosomes were grouped as efficiently translated mRNAs. A Student’s t-test was used to determine statistical significance (* p<0.05, ** p<0.005, *** p<0.0005). (**G**) Quantitative real-time PCR analysis of *CD155(PVR)* in human H441 cells treated for 24 hours with 100uM or 200uM Salubrinal or DMSO vehicle control and Actinomycin D (10ug/ml, 0 to 8 hours). *CD155* mRNA was normalized to *Actin*. (**H**) Western blot analysis of human H441 cells treated for 24 hours with 100uM Salubrinal or DMSO vehicle control and Cycloheximide (40ug/ml, 0 to 24 hours). CD155 protein quantification was normalized to vinculin.

**Supplementary Figure 3: The effect of ISR pathway activation on dendritic cells and T cells.** (**A**) ELISAs for IL-2 and Granzyme B of Jurkat T cells co-cultured with H358 cells cultured in RPMI 1640 media supplemented with or without amino acids and 800nM ISRIB, then washed and co-cultured with Jurkat T cells for an additional 24 hours with α-CD3 and α-CD28 activating antibodies (4ug/ml each). Error bars represent standard deviation from the mean from n=3 biological replicates. Data from a single experiment are shown and representative of three independent experiments. A Student’s t-test was used to determine statistical significance (** p<0.005). (**B**) Western blot analysis of murine JAWS II dendritic cells treated for 24 hours with 100uM or 200uM Salubrinal or DMSO vehicle control. (**C**) Western blot analysis of primary murine BMDCs (bone marrow dendritic cells) treated for 24 hours with 100uM Salubrinal, 5uM thapsigargin, 5-100 uM Arsenite, or DMSO vehicle control. (**D**) Western blot analysis of human Jurkat T cells treated for 24 hours with 5uM thapsigargin, in RPMI 1640 media supplemented with or without amino acids, or for 24 or 48 hours with 100uM or 200uM Salubrinal or DMSO vehicle control. Data from a single experiment are shown and representative of at least 3 independent experiments.

**Supplementary Figure 4: The effect of ISR pathway inhibition *in vitro* and *in vivo***. (**A**) Western blot analysis of *KRAS* mutant murine CMT167 cells treated for 24 hours with 5uM thapsigargin or DMSO vehicle control and 500nM ISRIB. (**B,C**) Quantification of tumor volumes of CMT167 cells expressing the indicated shRNA sequence transplanted in C57BL/6J mice (n= 15 mice per group). Graph represents mean tumor volumes and error bars represent standard error of the mean. (**D**) Diagram of gating strategy to gate live, single cells from flow mass cytometry (mass CyTOF). Cells were gated by event length/time, center/time, offset/time, width/time, and residual/time. Gating was used to remove signal from beads, and then further gating was performed to isolate live cells. (**E**) Gating strategy for TIL staining of cells from tumors. Cells were gated Cisplatin-/CD45+ (live, CD45+ TILs), then further gated based on TIL subtype. (**F**) CD8+ T cells were gated as in (**E**), CD45+ live cells were further gated CD3+, CD8+ to obtain %CD8+ (of CD45+ cells).

**Supplementary Figure 5: The effect of ISR pathway inhibition on immune cell populations *in vivo***. (**A**) Uniform manifold approximation and projection (UMAP) analysis of tumor infiltrating lymphocytes (TILs) colored by cell types. (**B**) Quantification of TILs (expressed as a % of CD45^+^ cells). n=5 *scrambled* mice, n=5 *Urod* shRNA mice, n=5 *Urod* shRNA + ISRIB, n=4 *Urod* shRNA + αPD-1, n=5 *Urod* shRNA + ISRIB +αPD-1. Graphs represent mean values and error bars represent standard deviation from the mean. A Student’s t-test was used to determine statistical significance (* p<0.05, ** p<0.005, *** p<0.0005).

**Supplementary Figure 6: Characterization of CD155 expression in surgically resected primary NSCLC patients.** (**A**) Microphotographs of CD155 immunohistochemistry analysis of primary lung adenocarcinomas displaying different levels of expression: negative, low, moderate and strong protein membrane expression in malignant cells. 20X representative images shown, scale bar=100μm (**B**) Association of CD155 H-score expression in all primary NSCLC with PD- L1 status (cutoff for positive expression Tumor proportion score >=1%) (p = 0.0005). A Mann Whitney test was used to determine statistical significance. (**C**) Association of CD155 H-score expression and % of CD155+ malignant cells with tumor histological diagnosis. (p<0.005). A Mann Whitney test was used to determine statistical significance. (**D**) Association of CD155 H-score expression with patient’s sex in all primary NSCLC and all primary lung adenocarcinoma. (* p<0.05, *** p<0.0005). A Mann Whitney test was used to determine statistical significance. (**E**) Association of CD155 H-score expression with patient’s smoking status in all primary NSCLC and all primary lung adenocarcinoma. Tukey’s multiple comparisons test was used to determine statistical significance. (* p<0.05, **** p<0.0001). (**F**) Association of CD155 H-score expression in all primary NSCLC with pathological stage. A Mann Whitney test was used to determine statistical significance (p<0.0001).

## Supplementary Tables

**Supplementary Table 1.** Clinicopathological characteristics of surgically resected primary NSCLC patients included in this study.

**Supplementary Table 2.** Chemicals used in this study.

**Supplementary Table 3.** Primers used in this study.

**Supplementary Table 4.** Antibodies used in this study.

