## Supplementary Figures and Tables for "Coordinated translational control of multiple immune checkpoints by the integrated stress response pathway in lung cancer"

### Supplementary Figure 1

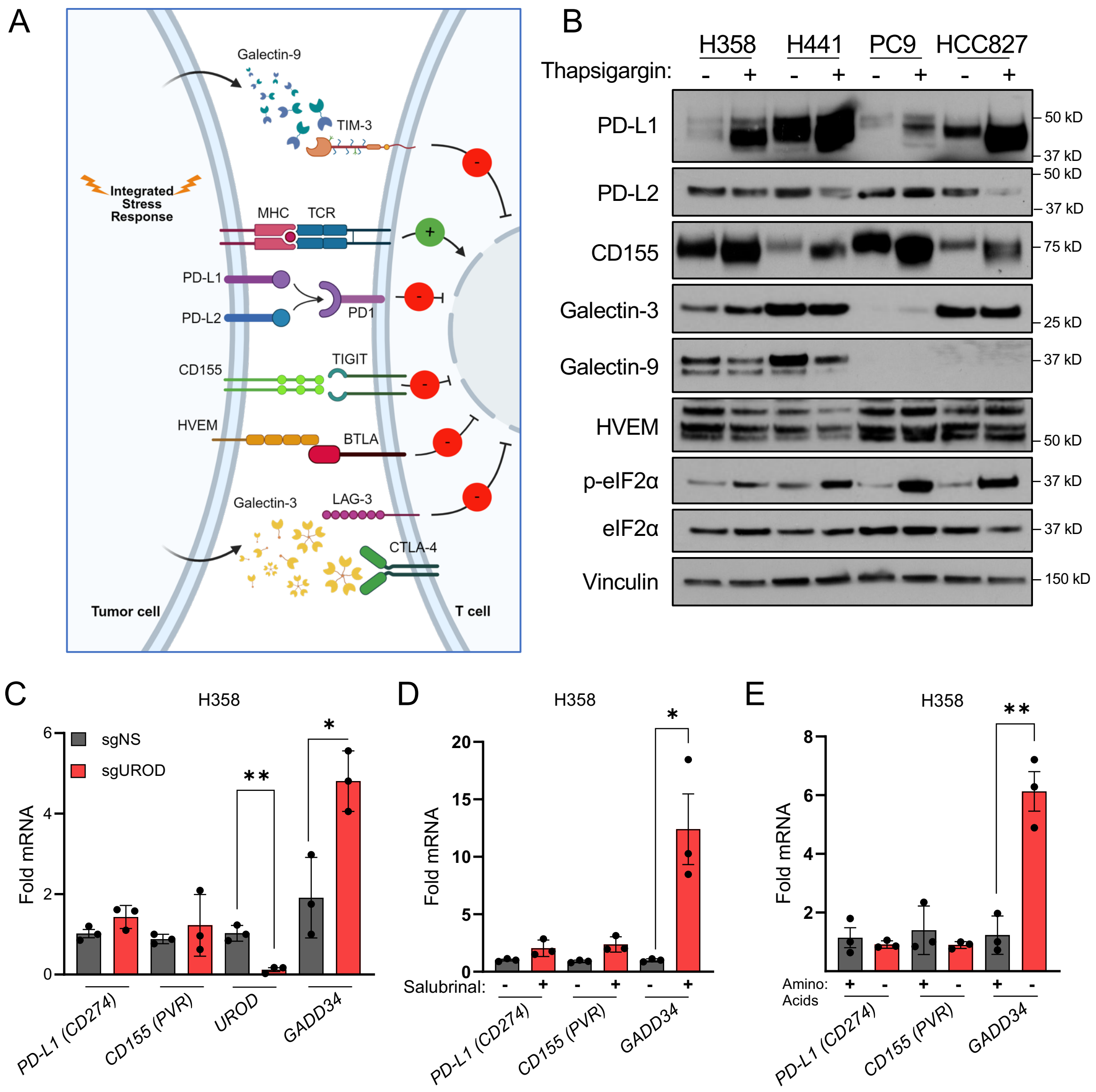

### Supplementary Figure 2

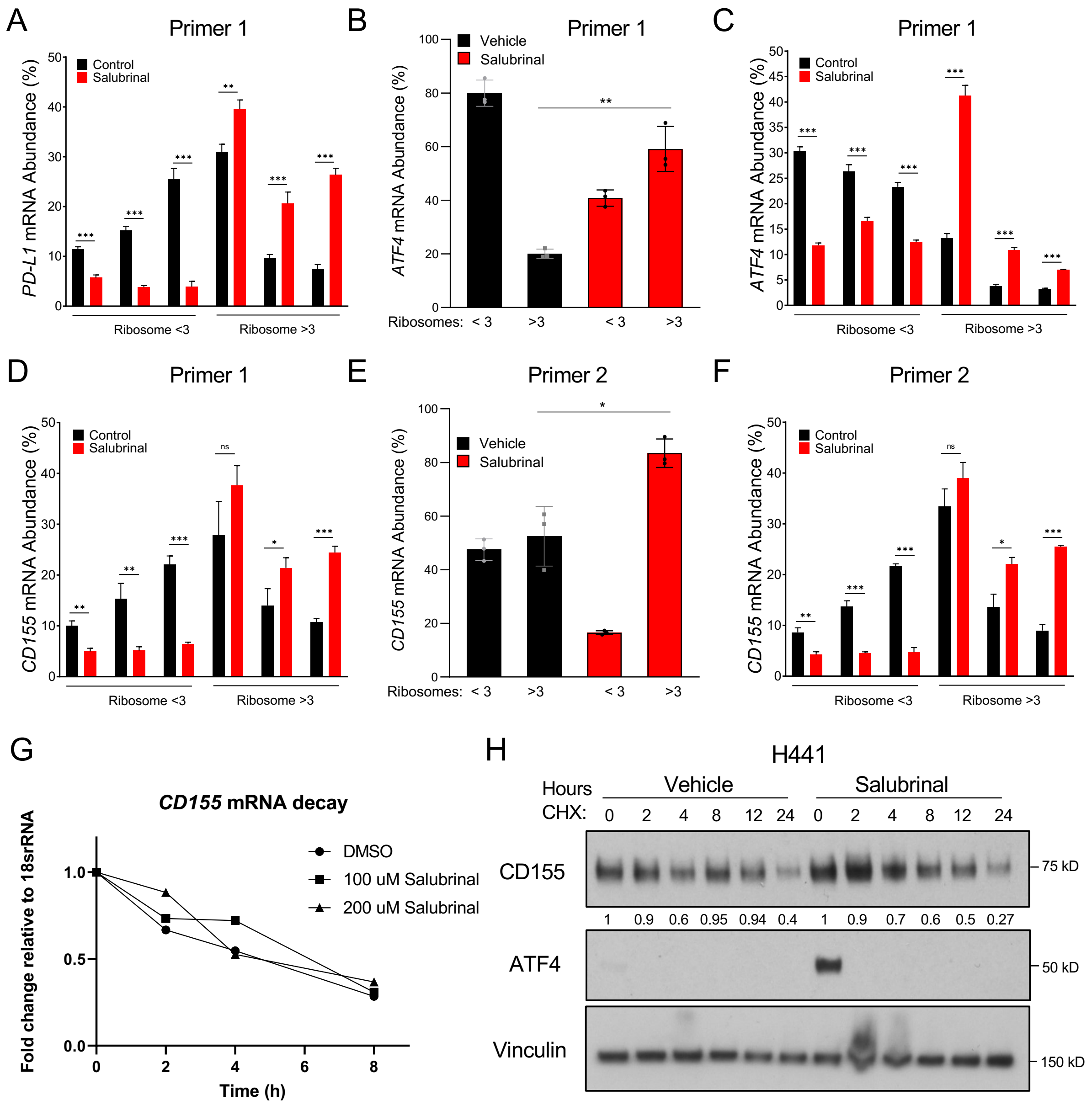

#### Supplementary Figure 3

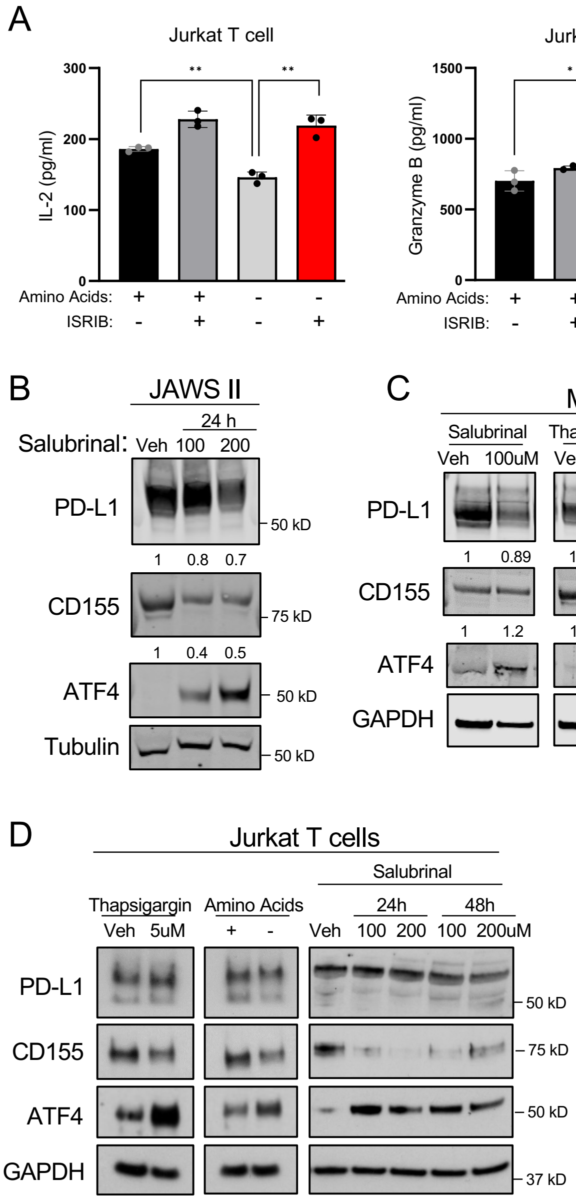

### Supplementary Figure 4

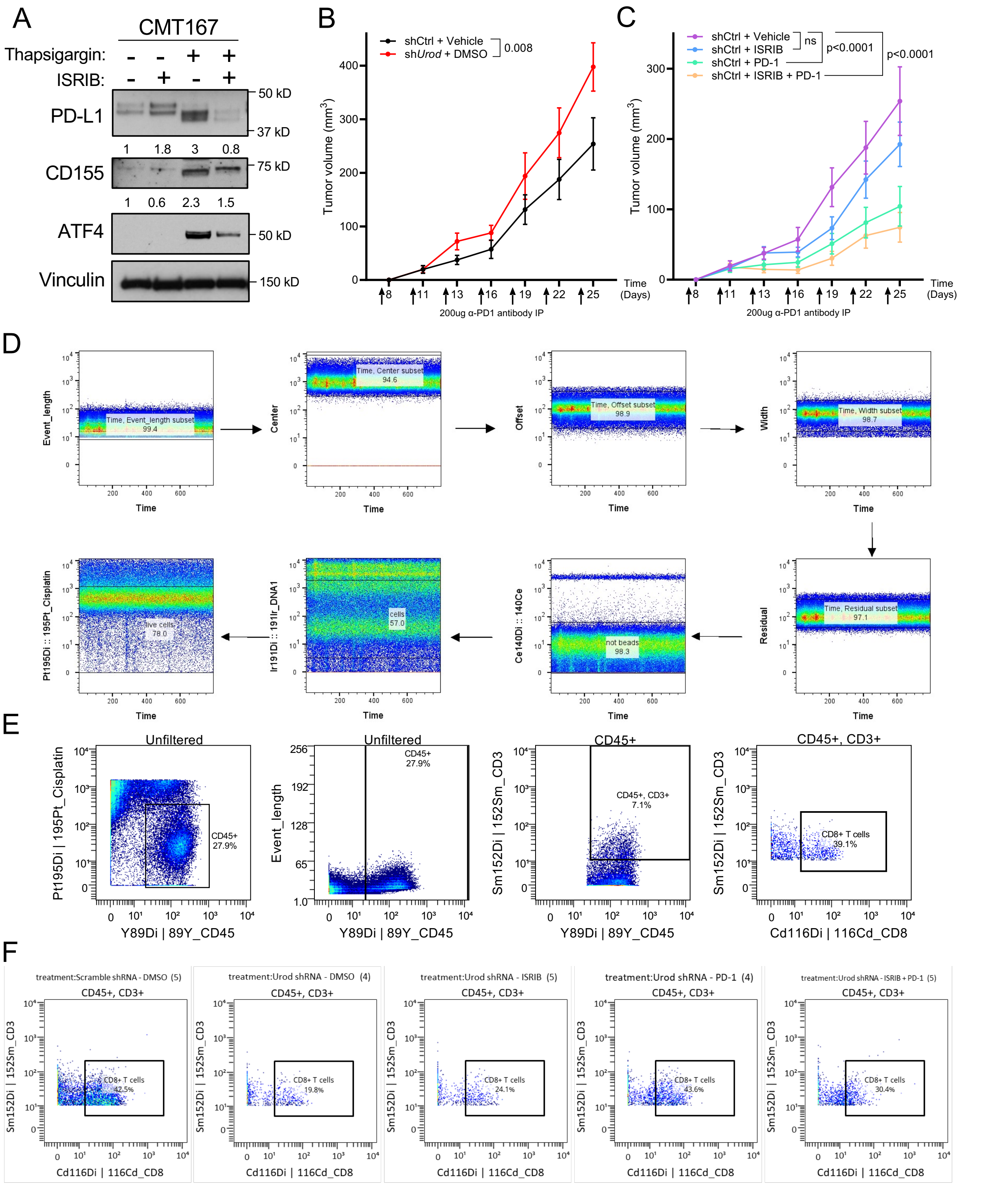

### Supplementary Figure 5

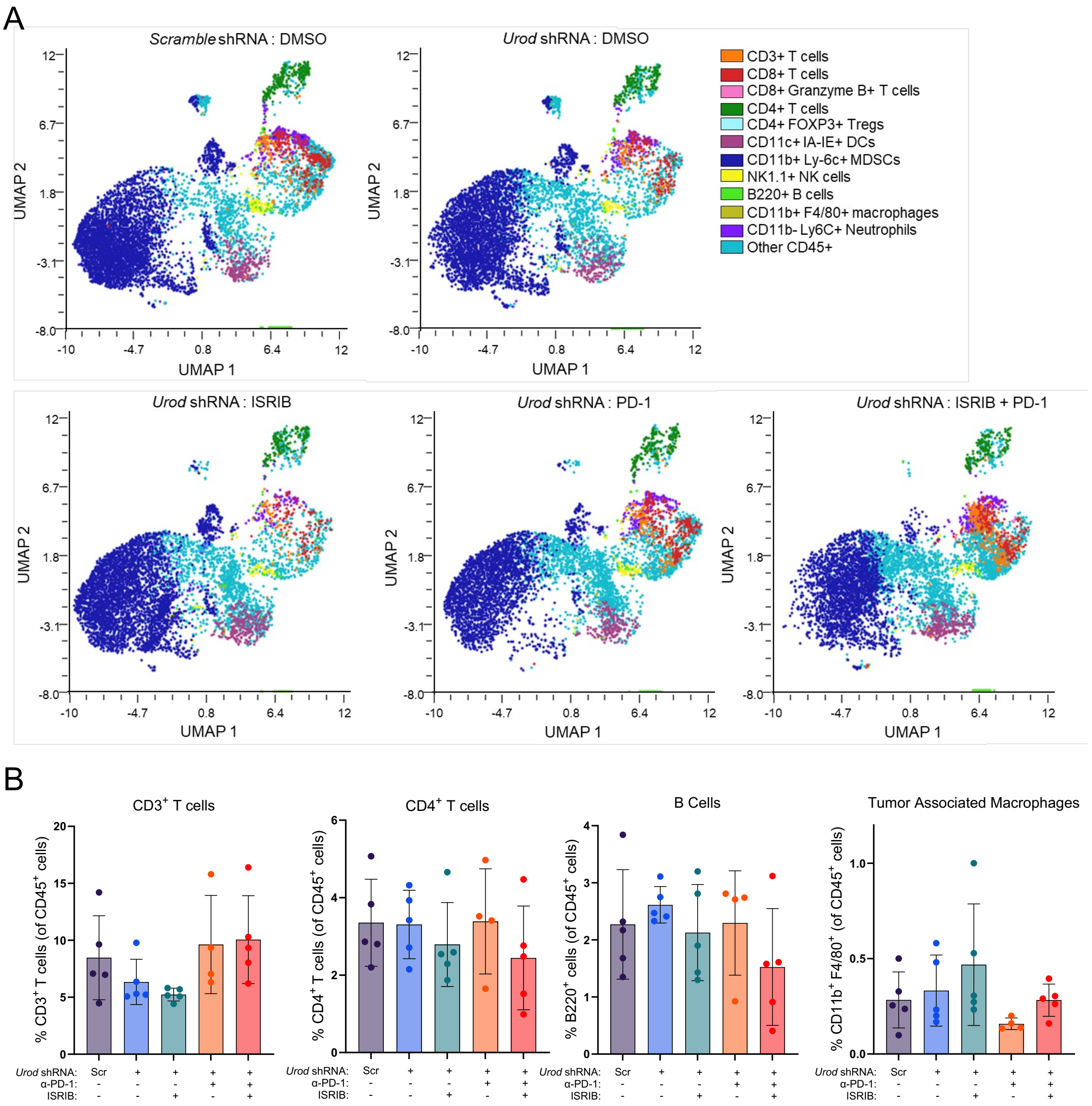

### Supplementary Figure 6

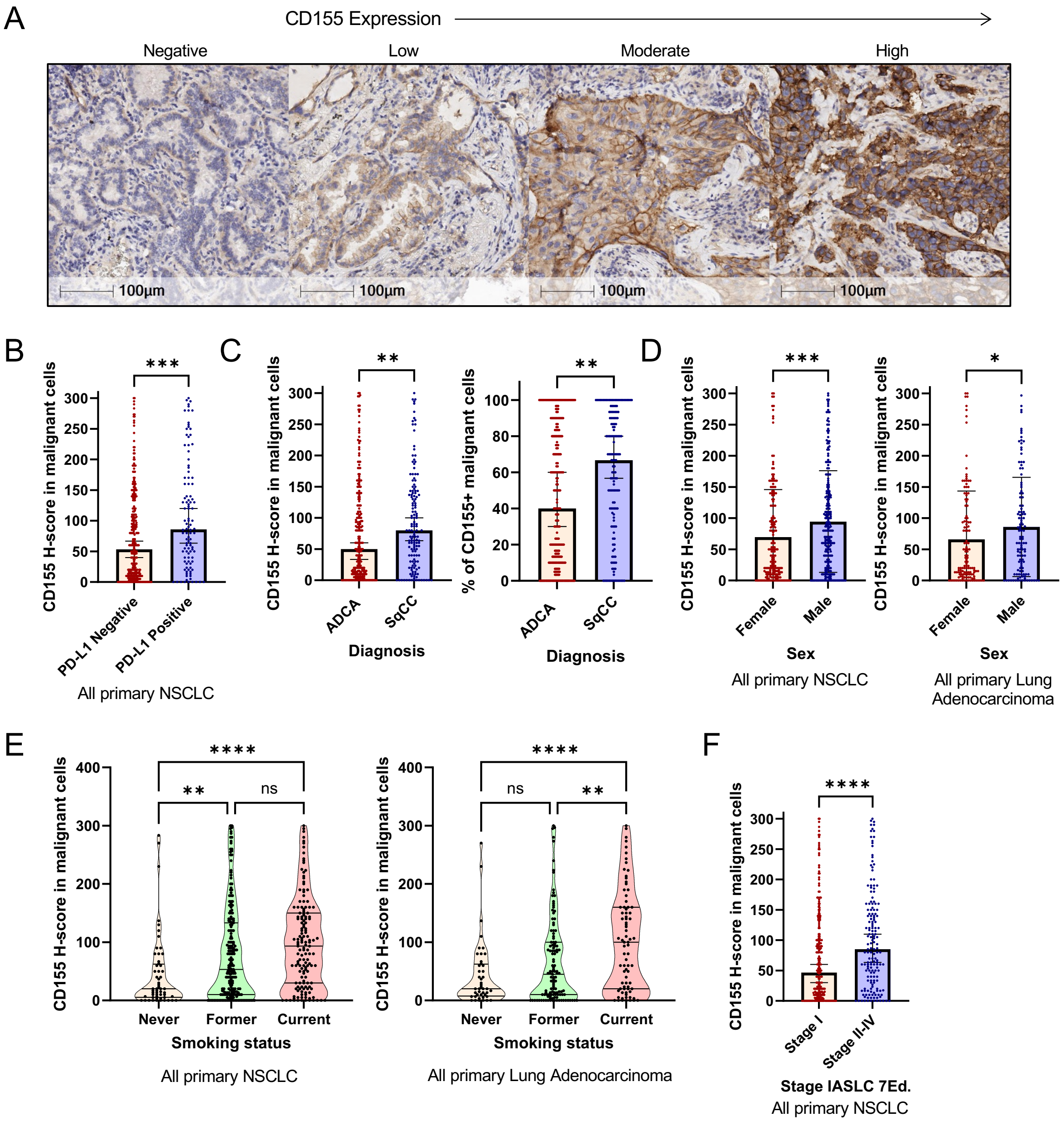

**Supplementary Table 1.**
**Clinicopathological characteristics of surgically resected primary NSCLC patients included in this study (N=410).**

| Features | | CD155 membrane IHC | | | | | |
|---|---|---|---|---|---|---|---|
| | | Negative (<1%) (n=52) | | Positive (>=1%) (n=358) | | Total (n=410) | P value |
| | | N | % | N | % | Total | |
| **Sex** | | | | | | | |
| | Female | 25 | 14 | 159 | 86 | 184 | ns |
| | Male | 27 | 12 | 199 | 88 | 226 | |
| **Smoking Status** | | | | | | | |
| | Never | 10 | 18 | 46 | 82 | 56 | ns |
| | Former | 30 | 14 | 177 | 86 | 207 | |
| | Current | 12 | 8 | 135 | 92 | 147 | |
| **Histolo** | | | | | | | |
| | Adenocarcinoma | 35 | 14 | 212 | 86 | 247 | ns |
| | Squamous cell carcinoma | 15 | 10 | 130 | 90 | 145 | |
| | Other NSCLC | 2 | 11 | 16 | 89 | 18 | |
| **Stage IASLC 7th edition** | | | | | | | |
| | I | 40 | 16 | 206 | 84 | 246 | 0.0223 |
| | II | 5 | 5 | 93 | 95 | 98 | Fisher's exact test |
| | III | 6 | 10 | 54 | 90 | 60 | |
| | IV | 1 | 17 | 5 | 83 | 6 | |
| **Stage IASLC 7th edition** | | | | | | | 0.0077 |
| | I | 40 | 16 | 206 | 84 | 246 | Chi-square test |
| | II-IV | 12 | 7 | 152 | 93 | 164 | |

**Supplementary Table 2. Chemicals used in this study.**

| Reagents | Source | Catalog # |
| --- | --- | --- |
| Salubrinal | Tocris | 23-471-0 |
| ISRIB | Fisher Scientific | 52-845-0 |
| Thapsigargin | Millipore Sigma | T9033-1MG |
| Cycloheximide | Millipore Sigma | C7698-5G |
| Actinomycin D | Thermo Fisher | A7592 |
| Blasticidin | Thermo Fisher | R21001 |
| Puromycin | Gibco | A1113803 |
| Doxycycline | RPI | D43020-250.0 |
| RNAseOUT | Millipore Sigma | R2020-250ML |
| Protease Inhibitor Cocktail, EDTA-Free | Fisher Scientific | PI87785 |
| Invitrogen™ Buffer Kit, RNase-free | Fisher Scientific | AM9010 |
| HEPES buffer | Fisher Scientific | NC1584172 |
| RNasin® Ribonuclease Inhibitor | Promega | N2515 |
| Glycogen | Roche | 10901393001 |
| Sucrose | Fisher Scientific | 50-188-2396 |
| DNAse I | Millipore Sigma | 10104159001 |
| Collagenase | Millipore Sigma | 10103586001 |
| (Hydroxypropyl)methyl cellulose | Millipore Sigma | H7509-25G |
| Tween 20 | Millipore Sigma | P9416-100ML |
| Triton X-100 | Millipore Sigma | T8787-50ML |

**Supplementary Table 3. Primers used in this study.**

| qPCR Primers | Sequence (5' to 3') |
|---|---|
| UROD qPCR FP | AGGCCTGCTGTGAACTGACT |
| UROD qPCR RP | CCTGGGGTACAACAAGGATG |
| hActin- forward | GCACAGAGCCTCGCCTTT |
| hActin- reverse | TATCATCATCCATGGTGAGCTGG |
| hPDL1-1 reverse | TGGTAATTCTGGGAGCCATC |
| hPDL1-1 forward | TCTTTGAGTTTGTATCTTGGATGC |
| hPDL1-2 forward | GCTGCATGATCAGCTATGGT |
| hPDL1-2 reverse | GTGACTGGATCCACAACCAA |
| hCD155 forward | GTCCAAATGTTCCCGTGAGG |
| hCD155 reverse | GCTGAATAGGAGACATGCCCA |
| hCD155-2 forward | TTTGGCACTGTCATCTGTGTC |
| hCD155-2 reverse | GCTGTCACCTTGTGCCCTC |
| hGADD34 forward | AAGGCCAGAAAGGTGCGCTT |
| hGADD34 reverse | GCGATCCCGAGCAAGCTG |
| Luciferase forward | GAGGCGAACTGTGTGTGAGA |
| Luciferase reverse | GAGCCACCTGATAGCCTTTG |

**Supplementary Table 4. Antibodies used in this study.**

| Antigen | Source | Catalog # | Application |
|---|---|---|---|
| Vinculin | Abcam | ab129002 | WB |
| PD-L1 | CST | 13684S | WB, IHC |
| CD155 | CST | 81254S | WB, IHC |
| p-eIF2α | Abcam | ab32157 | WB |
| eIF2α | CST | 9722S | WB |
| eIF5B | Santa Cruz | sc-393564 | WB |
| UROD (UPD) | Abcam | ab196562 | WB |
| eIF3D | Bethyl | 4301-758A | WB |
| eIF3A | CST | 3411S | WB |
| eIF4E | CST | 2067S | WB |
| eIF6 | CST | 3833S | WB |
| Galectin-3 | CST | 87985S | WB |
| Galectin-9 | Biorad | VMA00212 | WB |
| PD-L2 | Abcam | ab256386 | WB |
| HVEM | Santa Cruz | sc-365971 | WB |
| GAPDH | CST | 5174S | WB |
| Mouse-specific PD-L1 | Abcam | ab213480 | WB |
| Mouse-specific CD155 | Thermo-Fisher | MA5-29762 | WB |
| APC-PD-L1 | Biolegend | 329708 | Flow cytomery |
| FITC-CD155 | Biolegend | 337628 | Flow cytometry |
| CD8α | CST | 98941S | IHC-F |
| CD4 | CST | 25229S | IHC-F |
| CD3ε | CST | 78588S | IHC-F |
| CD3, Functional Grade, eBioscience | Thermo Fisher | 16-0037-85 | *in vitro* |
| CD28, Functional Grade, eBioscience | Thermo Fisher | 16-0289-85 | *in vitro* |
| *InvivoMab* anti-mouse PD-1 antibody clone 29F.1A12 | BioXCell | BE0273 | *in vivo* |
| *InvivoMab* rat IgG2a isotype control clone 2A3 | BioXCell | BE0089 | *in vivo* |
| CD45 89Y | Fluidigm | 3089005B | Mass CyTOF |
| CD45/B220 176Yb | Fluidigm | 3176002B | Mass CyTOF |
| CD8 116CD | Biolegend | 100755 | Mass CyTOF |
| CD3e 152Sm | Fluidigm | 3152004B | Mass CyTOF |
| CD4 145Nd | Fluidigm | 3145002B | Mass CyTOF |
| CD11b 148Nd | Fluidigm | 3148003B | Mass CyTOF |
| CD11c 142Nd | Fluidigm | 3142003B | Mass CyTOF |
| CD19 149Sm | Fluidigm | 3149002B | Mass CyTOF |
| CD25 151Eu | Fluidigm | 3151007B | Mass CyTOF |
| CD206 175Lu | Biolegend | 151702 | Mass CyTOF |
| IA/IE 209Bi | Fluidigm | 3209006B | Mass CyTOF |
| Ly-6C 162Dy | Fluidigm | 3162014B | Mass CyTOF |
| Ly-6G 141Pr | Fluidigm | 3141008B | Mass CyTOF |

| | | | |
|---|---|---|---|
| NK1.1 170ER | Fluidigm | 3170002B | Mass CyTOF |
| F4/80 146Nd | Fluidigm | 3146008B | Mass CyTOF |
| Ki-67 168Er | Fluidigm | 3168007B | Mass CyTOF |
| Foxp3 158Gd | CST | 3158003A | Mass CyTOF |
| PD-1 159Tb | Fluidigm | 3159024B | Mass CyTOF |
| PDL-1 153Eu | Fluidigm | 3153016B | Mass CyTOF |
| Perforin 172Yb | Fluidigm | 3172018B | Mass CyTOF |
| IFNy 165Ho | Fluidigm | 3165003B | Mass CyTOF |
| IL-2 144Nd | Fluidigm | 3144002C | Mass CyTOF |
| Granzyme B 171Yb | Fluidigm | 3171002B | Mass CyTOF |

CST = Cell Signaling Technologies
